# Social aging trajectories are sex-specific, sensitive to adolescent stress, and most robustly revealed during social tests with familiar stimuli

**DOI:** 10.1101/2023.04.27.538622

**Authors:** Christopher Figueroa, Erin L. Edgar, J. M. Kirkland, Ishan Patel, David N. King’uyu, Ashley M. Kopec

**Affiliations:** Department of Neuroscience and Experimental Therapeutics Albany Medical College

## Abstract

Social networks and support are integral to health and wellness across the lifespan, and social engagement may be particularly important during aging. However, social behavior and social cognition decline naturally during aging across species. Social behaviors are in part supported by the ‘reward’ circuitry, a network of brain regions that develops during adolescence. We published that male and female rats undergo adolescent social development during sex-specific periods, pre-early adolescence in females and early-mid adolescence males. Although males and females have highly dimorphic development, expression, and valuation of social behaviors, there is relatively little data indicating whether social aging is the same or different between the sexes. Thus, we sought to test two hypotheses: (1) natural social aging will be sex-speciifc, and (2) social isolation stress restricted to sex-specific adolescent critical periods for social development would impact social aging in sex-specific ways. To do this, we bred male and female rats in-house, and divided them randomly to receive either social isolation for one week during each sex’s respective critical period, or no manipulation. We followed their social aging trajectory with a battery of five tests at 3, 7, and 11 months of age. We observed clear social aging signatures in all tests administered, but sex differences in natural social aging were most robustly observed when a familiar social stimulus was included in the test. We also observed that adolescent isolation did impact social behavior, in both age-independent and age-dependent ways, that were entirely sex-specific. Please note, this preprint will not be pushed further to publication (by me, AMK), as I am leaving academia. So, it’s going to be written more conversationally.

## INTRODUCTION

Social networks and support are integral to health and wellness across the lifespan. Social engagement may be particularly important during aging, as social isolation in rodents shortens lifespan [1], perceived social isolation (i.e. loneliness) can increase mortality by up to 50% in humans [2, 3], and social stress exacerbates disease pathology in rodent models [4]. Conversely, frequent social activities in elderly individuals can reduce global cognitive decline by up to 70% [5], and strong social support promotes a variety of positive health outcomes in humans and rodent models [6–13]. However, social behavior and social cognition decline naturally during aging across species [14–18]. Moreover, although males and females have highly dimorphic expression and valuation of social behaviors [19–21], there is relatively little data indicating whether social aging is the same or different between the sexes. A better understanding of the best ways to assess social aging, and the factors that impact social aging for each sex will be important for preserving social function, and in turn health in a broad sense, in advanced age.

Social behaviors are in part supported by the ‘reward’ circuitry, a network of brain regions that develops during adolescence [22–24]. Interestingly, a large cross-sectional brain imaging study in humans identified brain activity networks that developed during adolescence that also exhibited accelerated degeneration in old age relative to the rest of the brain [25]. Prolonged social stress during adolescence induces a variety of reward circuitry and reward-related behavioral changes [26], and is reported to worsen spatial memory impairment in aged mice [27]. However, we published that male and female rats undergo adolescent social development during sex-specific periods, pre-early adolescence (postnatal day (P)20-30) in females and early-mid adolescence (P30-40) in males [28]. Disrupting natural developmental signaling within the reward circuitry between P20-30 in females, or between P30-40 in males, was sufficient to impair social development [28]. These data suggest that disrupting adolescent development will have sex-specific effects depending on *when during adolescence* the disruption occurs, a notion well-supported by the adolescent drug exposure literature [29, 30].

Thus, we sought to test two hypotheses: (1) natural social aging will be sex-speciifc, and (2) social isolation stress restricted to sex-specific adolescent critical periods for social development would impact social aging in sex-specific ways. To do this we bred male and female rats in-house, and divided them randomly to receive either social isolation for one week during each sex’s respective critical period, or no manipulation. We followed their social aging trajectory with a battery of five tests at 3, 7, and 11 months (mo) of age. We observed clear social aging signatures in all tests administered, but sex differences in natural social aging were most robustly observed when a familiar social stimulus was included in the test. We also observed that adolescent isolation did impact social behavior, in both age-independent and age-dependent ways, that were entirely sex-specific.

## METHODS

### Animal Care

Adult male and female Sprague-Dawley rats were purchased to be breeding pairs (Harlan/Envigo). Litters were culled to a maximum of 12 pups between postnatal day (P)2-5, and pups were weaned into same-sex pair-housing at P21. No more than two rats per sex per litter were used in each condition. For Juvenile Social Interaction and Social Choice tests, rats were purchased from Harlan/Envigo to act as conspecifics. Rats were housed in conventional cages on cellulose bedding with ad libitum access to food and water. Colonies were maintained on a 12:12 light:dark cycle (lights on a 07:00) in a temperature-(20-24°C) and humidity-(35-55 RH) controlled room. Cages were changed twice weekly until ∼9mo of age, when they were changed thee times per week. All experiments were approved by the Institutional Animal Care and Use Committee at Albany Medical College.

### Animal Model

Female rats were single-housed (social isolation) from P23-29 and male rats were single-housed from P31-37, all of which were re-pair-housed with their previous cage mates after the manipulation. Control male and female rats remained in pair-housing throughout the studies. Cage mates always received the same manipulation. Behavior was tested at 3 months, 7-8 months (hereafter referred to as 7 months), and 11-12 months (hereafter referred to as 11 months). Prior to behavioral testing, both experimental and conspecific rats were handled 3 times for ∼5mins in the behavior room. Experimental rats were also acclimated for 10 mins in the Choice test arenas once prior to testing at each age. Five total tests were conducted at each age, once daily from Mon – Fri, between 08:00-11:00 and 15:00-18:00 to minimize sleep cycle disruption. Behavior test order was counterbalanced. All rats were euthanized and tissues collected at ∼13mo of age (the conclusion of the study).

### Behavioral Testing

#### Social vs. Object Choice test

The choice test arena consists of two chambers designated the social or nonsocial halves. Flanking each chamber is an enclosed space for holding the stimuli that is separated from the exploration area by a clear plexiglass partition with holes to permit visual and olfactory exploration, but no physical contact. Rats were acclimated to the behavior room for ∼20 mins on test day, and had been previously acclimated to experimenter handling and the Choice test arena. A sex-matched novel adult conspecific rat (≥P90) was placed in the social restriction space and a beaker of marbles was placed in the nonsocial enclosure space (**Supp. Fig. 1A**).

Stimuli placement was counterbalanced. Experimental rats were placed in the exploration area for 10 mins and recorded using AnyMaze software from overhead cameras. Time spent exploring the stimuli and in each half of the arena were hand-coded by a blinded experimenter using Solomon Coder (András Péter). We experimentally determined that 5 mins of coding is sufficient to capture the behavioral trends of the experimental rats (*data not shown*), and thus we coded just the first 5 mins of the test. Following the test, rats were replaced in their home cages.

#### Social Novelty Preference test

The same arena and procedures described in Social vs. Object Choice was used for Social Novelty Preference tests. The two chambers were designated as novel social or familiar social halves. A sex-matched novel adult conspecific rat (≥P90) was placed in the novel social restriction space and the experimental rats’ familiar cage mate was placed in the familiar social enclosure space (**Supp. Fig. 1B**). Stimuli placement was counterbalanced.

#### Familiar Interaction

Cage mates were separated for 20 mins while each was acclimated to a new, clean rat cage with no food/water hopper or enrichment for 10 mins each. Cage mates were paired together in the clean cage for 10 mins and recorded using AnyMaze software and parallel cameras. Videos were hand-coded by a blinded experimenter for (**Supp. Fig. 1C**):

a. active social interaction: the experimental rat is visually attending to and physically in contact with its partner
b. passive social interaction: the experimental rat is visually attending to, but not physically in contact with, its partner
c. nonsocial contact: the experimental rat is not visually attend to, but is in physical contact with, its partner
d. nonsocial attention: the experimental rat is neither visually attending to nor physically in contact with its partner

Each video was coded twice, once for each cage mate. We experimentally determined that 5 mins of coding is sufficient to capture the behavioral trends of the rats (*data not shown*), and thus we coded just the first 5 mins of the test.

#### Novel Juvenile Interaction

Experimental rats were acclimated to a new, clean rat cage with no food/water hopper or enrichment for 10 mins. A sex-matched juvenile rat (P25-27) was introduced to the cage for 10 mins and video and coding was completed by a blinded experimenter as described in Familiar Interaction. Each video was coded once, just for the experimental rat.

### Data Analysis and Statistics

Data analyzed from Choice tests include (1) total exploration of each stimulus and (2) time spent exploring the stimulus as a percentage of the total time spent in the corresponding half of the arena. Data from the Interaction tests include the four social measurements described above. Because of modest group sizes, we did not test for outliers or exclude any data. We analyzed both sex differences in natural social aging (control males and females) and the effect of adolescent manipulation on social aging (control males vs. P31-37 isolation males and control females vs. P23-29 isolation females). Rats were assessed on each test at 3 different ages, resulting in mixed model two-way ANOVAs (between: sex or adolescent manipulation, within: age). Significant main effects or interactions were followed by post-hoc tests. Statistical significance is defined as *p*<0.05. All statistics were performed in GraphPad Prism 9.5.1.

## RESULTS

Our experimental design required separate analysis of sex differences in natural aging and the effect of adolescent isolation for each sex, as the critical periods for isolation were sex-specific and defined by our previous work [28]. In this Results section, we will present the main conclusions for each set of analyses for ease of reading, but full statistical details can be found in **Tables 1-5**.

**Table 1:** Statistical details for Social vs. Object choice test

|  | Measurement | ANOVA Factor | Statistic | p-value | Post-hoc | Figure |
| --- | --- | --- | --- | --- | --- | --- |
| Sex Differences in Natural Aging | Social Exploration | Age | F(2,32)=8.82 | 0.001 | 3mo > 7mo & 11mo | 1A |
|  |  | Sex | F(1,16)=1.02 | 0.327 |  |  |
|  |  | Age x Sex | F(2,32)=0.22 | 0.806 |  |  |
|  | Object Exploration | Age | F(2,32)=6.41 | 0.005 | 11mo > 3mo & 7mo | 1B |
|  |  | Sex | F(1,16)=10.07 | 0.006 | Female > Male |  |
|  |  | Age x Sex | F(2,32)=0.11 | 0.894 |  |  |
|  | % Social Exploration | Age | F(2,32)=3.95 | 0.029 | 3mo > 11mo | 1C |
|  |  | Sex | F(1,16)=5.96 | 0.027 | Female > Male |  |
|  |  | Age x Sex | F(2,32)=0.01 | 0.987 |  |  |
|  | % Object Exploration | Age | F(2,32)=4.14 | 0.025 | 11mo > 7mo | 1D |
|  |  | Sex | F(1,16)=0.09 | 0.773 |  |  |
|  |  | Age x Sex | F(2,32)=0.18 | 0.836 |  |  |
| Effect of Adolescent Isolation in Males | Social Exploration | Age | F(2,30)=6.97 | 0.003 | 3mo > 7mo & 11mo | 6A |
|  |  | Manipulation | F(1,16)=0.80 | 0.383 |  |  |
|  |  | Age x Manipulation | F(2,30)=0.08 | 0.924 |  |  |
|  | Object Exploration | Age | F(2,46)=9.1 | 0.001 | 11mo > 3mo & 7mo | 6B |
|  |  | Manipulation | F(1,46)=0.67 | 0.416 |  |  |
|  |  | Age x Manipulation | F(2,46)=0.01 | 0.993 |  |  |
|  | % Social Exploration | Age | F(2,30)=4.54 | 0.019 | 3mo > 7mo | 6C |
|  |  | Manipulation | F(1,16)=0.85 | 0.370 |  |  |
|  |  | Age x Manipulation | F(2,30)=0.57 | 0.569 |  |  |
|  | % Object Exploration | Age | F(2,46)=2.32 | 0.110 |  | 6D |
|  |  | Manipulation | F(1,46)=4.86 | 0.033 | Ctrl > P31-37iso |  |
|  |  | Age x Manipulation | F(2,46)=1.07 | 0.350 |  |  |
| Effect of Adolescent Isolation in Females | Social Exploration | Age | F(2,24)=8.88 | 0.001 | 3mo > 7mo & 11mo | 6E |
|  |  | Manipulation | F(1,14)=0.01 | 0.928 |  |  |
|  |  | Age x Manipulation | F(2,24)=0.42 | 0.665 |  |  |
|  | Object Exploration | Age | F(2,24)=3.31 | 0.054 |  | 6F |
|  |  | Manipulation | F(1,14)=1.40 | 0.257 |  |  |
|  |  | Age x Manipulation | F(2,24)=0.12 | 0.892 |  |  |
|  | % Social Exploration | Age | F(2,24)=5.48 | 0.011 | 3mo > 7mo | 6G |
|  |  | Manipulation | F(1,14)=0.04 | 0.835 |  |  |
|  |  | Age x Manipulation | F(2,24)=0.73 | 0.493 |  |  |
|  | % Object Exploration | Age | F(2,38)=4.90 | 0.013 | 3mo & 11mo > 7mo | 6H |
|  |  | Manipulation | F(1,38)=5.66 | 0.023 | P23-29iso > Ctrl |  |
|  |  | Age x Manipulation | F(2,38)=0.25 | 0.783 |  |  |

**Table 2:** Statistical details for Social Novelty Preference choice test

|  | Measurement | ANOVA Factor | Statistic | p-value | Post-hoc | Figure |
| --- | --- | --- | --- | --- | --- | --- |
| Sex Differences in Natural Aging | Novel Exploration | Age | F(2,45)=6.75 | 0.003 | 3mo & 7mo > 11mo | 2A |
|  |  | Sex | F(1,45)=1.05 | 0.311 |  |  |
|  |  | Age x Sex | F(2,45)=3.55 | 0.037 | 3mo > 11mo in Males only |  |
|  | Familiar Exploration | Age | F(2,29) = 15.14 | <0.001 | 3mo > 7mo & 11mo | 2B |
|  |  | Sex | F(1,16)=4.13 | 0.059 |  |  |
|  |  | Age x Sex | F(2,29)=3.57 | 0.041 | 3mo > 7mo & 11mo in Females only |  |
|  | % Novel Exploration | Age | F(2,45)=17.15 | <0.001 | 3mo & 7mo > 11mo | 2C |
|  |  | Sex | F(1,45)=13.69 | 0.001 | Females > Males |  |
|  |  | Age x Sex | F(2,45)=0.37 | 0.693 |  |  |
|  | % Familiar Exploration | Age | F(2,45)=21.56 | <0.001 | All ages different | 2D |
|  |  | Sex | F(1,45)=1.17 | 0.285 |  |  |
|  |  | Age x Sex | F(2,45)=1.60 | 0.212 |  |  |
| Effect of Adolescent Isolation in Males | Novel Exploration | Age | F(2,30)=11.37 | <0.001 | 3mo & 7mo > 11mo | 7A |
|  |  | Manipulation | F(1,16)=3.47 | 0.081 |  |  |
|  |  | Age x Manipulation | F(2,30)=1.31 | 0.285 |  |  |
|  | Familiar Exploration | Age | F(2,30)=7.47 | 0.002 | 3mo > 7mo & 11mo | 7B |
|  |  | Manipulation | F(1,16)=0.01 | 0.924 |  |  |
|  |  | Age x Manipulation | F(2,30)=0.80 | 0.457 |  |  |
|  | % Novel Exploration | Age | F(2,30)=18.55 | <0.001 | 3mo & 7mo > 11mo | 7C |
|  |  | Manipulation | F(1,16)=3.08 | 0.099 |  |  |
|  |  | Age x Manipulation | F(2,30)=0.15 | 0.861 |  |  |
|  | % Familiar Exploration | Age | F(2,30)=13.59 | <0.001 | 3mo > 7mo & 11mo | 7D |
|  |  | Manipulation | F(1,16)=0.13 | 0.725 |  |  |
|  |  | Age x Manipulation | F(2,30)=0.05 | 0.954 |  |  |
| Effect of Adolescent Isolation in Females | Novel Exploration | Age | F(2,21)=6.73 | 0.006 | 3mo & 7mo > 11mo | 7E |
|  |  | Manipulation | F(1,14)=0.00 | 0.987 |  |  |
|  |  | Age x Manipulation | F(2,21)=3.17 | 0.063 |  |  |
|  | Familiar Exploration | Age | F(2,21)=12.14 | <0.001 | 3mo > 7mo & 11mo | 7F |
|  |  | Manipulation | F(1,14)=0.20 | 0.662 |  |  |
|  |  | Age x Manipulation | F(2,21)=4.71 | 0.020 | 3mo > 7mo & 11mo in Ctrl only |  |
|  | % Novel Exploration | Age | F(2,21)=10.67 | 0.001 | 3mo & 7mo > 11mo | 7G |
|  |  | Manipulation | F(1,14)=1.48 | 0.243 |  |  |
|  |  | Age x Manipulation | F(2,21)=0.21 | 0.816 |  |  |
|  | % Familiar Exploration | Age | F(2,21)=15.07 | <0.001 | 3mo & 7mo > 11mo | 7H |
|  |  | Manipulation | F(1,14)=0.38 | 0.549 |  |  |
|  |  | Age x Manipulation | F(2,21)=0.66 | 0.526 |  |  |

**Table 3:**
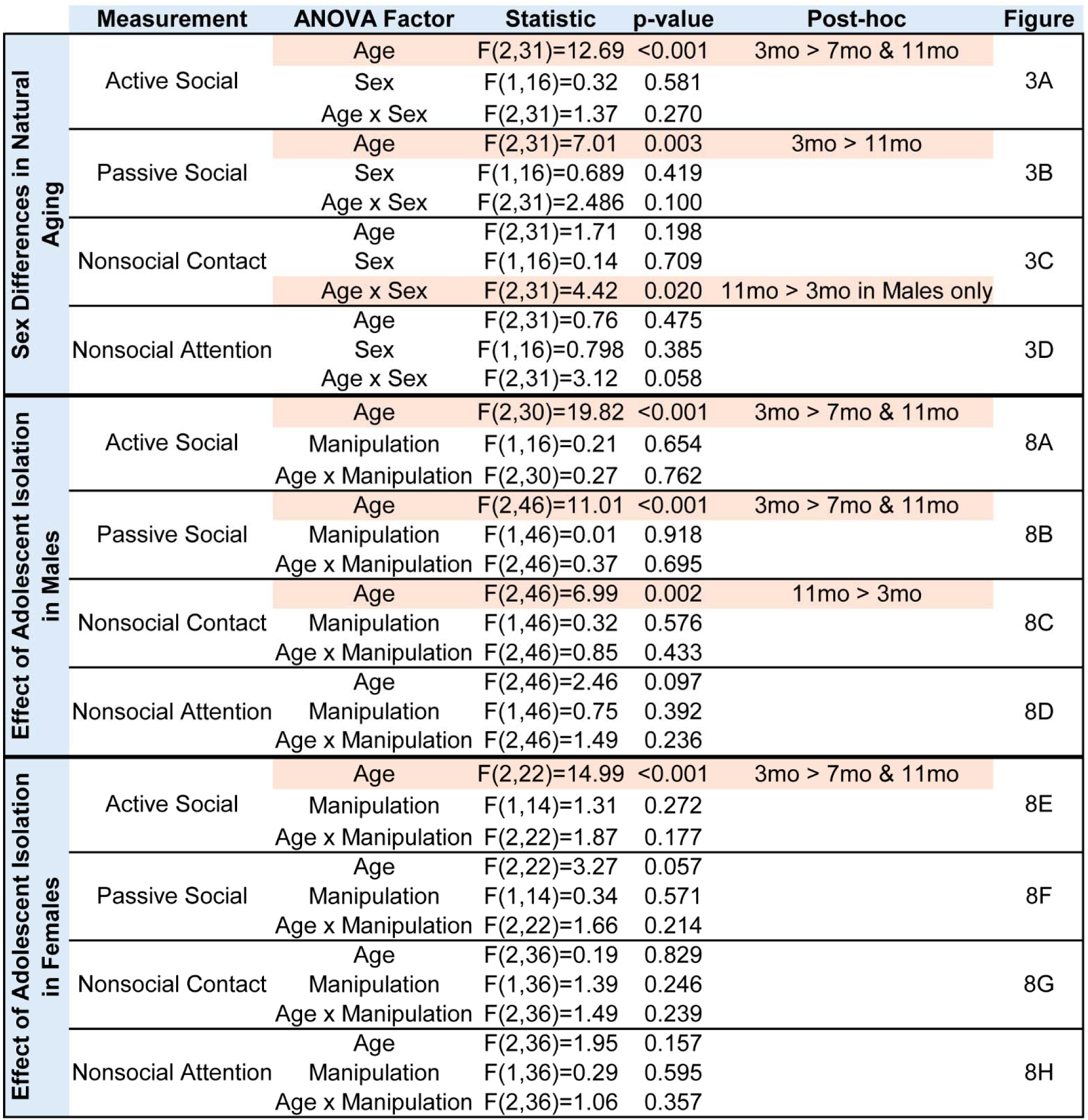
Statistical details for Novel Juvenile Social Interaction

|  | Measurement | ANOVA Factor | Statistic | p-value | Post-hoc | Figure |
| --- | --- | --- | --- | --- | --- | --- |
| Sex Differences in Natural Aging | Active Social | Age | F(2,31)=12.69 | <0.001 | 3mo > 7mo & 11mo | 3A |
|  |  | Sex | F(1,16)=0.32 | 0.581 |  |  |
|  |  | Age x Sex | F(2,31)=1.37 | 0.270 |  |  |
|  | Passive Social | Age | F(2,31)=7.01 | 0.003 | 3mo > 11mo | 3B |
|  |  | Sex | F(1,16)=0.689 | 0.419 |  |  |
|  |  | Age x Sex | F(2,31)=2.486 | 0.100 |  |  |
|  | Nonsocial Contact | Age | F(2,31)=1.71 | 0.198 |  | 3C |
|  |  | Sex | F(1,16)=0.14 | 0.709 |  |  |
|  |  | Age x Sex | F(2,31)=4.42 | 0.020 | 11mo > 3mo in Males only |  |
|  | Nonsocial Attention | Age | F(2,31)=0.76 | 0.475 |  | 3D |
|  |  | Sex | F(1,16)=0.798 | 0.385 |  |  |
|  |  | Age x Sex | F(2,31)=3.12 | 0.058 |  |  |
| Effect of Adolescent Isolation in Males | Active Social | Age | F(2,30)=19.82 | <0.001 | 3mo > 7mo & 11mo | 8A |
|  |  | Manipulation | F(1,16)=0.21 | 0.654 |  |  |
|  |  | Age x Manipulation | F(2,30)=0.27 | 0.762 |  |  |
|  | Passive Social | Age | F(2,46)=11.01 | <0.001 | 3mo > 7mo & 11mo | 8B |
|  |  | Manipulation | F(1,46)=0.01 | 0.918 |  |  |
|  |  | Age x Manipulation | F(2,46)=0.37 | 0.695 |  |  |
|  | Nonsocial Contact | Age | F(2,46)=6.99 | 0.002 | 11mo > 3mo | 8C |
|  |  | Manipulation | F(1,46)=0.32 | 0.576 |  |  |
|  |  | Age x Manipulation | F(2,46)=0.85 | 0.433 |  |  |
|  | Nonsocial Attention | Age | F(2,46)=2.46 | 0.097 |  | 8D |
|  |  | Manipulation | F(1,46)=0.75 | 0.392 |  |  |
|  |  | Age x Manipulation | F(2,46)=1.49 | 0.236 |  |  |
| Effect of Adolescent Isolation in Females | Active Social | Age | F(2,22)=14.99 | <0.001 | 3mo > 7mo & 11mo | 8E |
|  |  | Manipulation | F(1,14)=1.31 | 0.272 |  |  |
|  |  | Age x Manipulation | F(2,22)=1.87 | 0.177 |  |  |
|  | Passive Social | Age | F(2,22)=3.27 | 0.057 |  | 8F |
|  |  | Manipulation | F(1,14)=0.34 | 0.571 |  |  |
|  |  | Age x Manipulation | F(2,22)=1.66 | 0.214 |  |  |
|  | Nonsocial Contact | Age | F(2,36)=0.19 | 0.829 |  | 8G |
|  |  | Manipulation | F(1,36)=1.39 | 0.246 |  |  |
|  |  | Age x Manipulation | F(2,36)=1.49 | 0.239 |  |  |
|  | Nonsocial Attention | Age | F(2,36)=1.95 | 0.157 |  | 8H |
|  |  | Manipulation | F(1,36)=0.29 | 0.595 |  |  |
|  |  | Age x Manipulation | F(2,36)=1.06 | 0.357 |  |  |

**Table 4:** Statistical details for Familiar Cage mate Social Interaction

|  | Measurement | ANOVA Factor | Statistic | p-value | Post-hoc | Figure |
| --- | --- | --- | --- | --- | --- | --- |
| Sex Differences in Natural Aging | Active Social | Age | F(2,32)=3.89 | 0.031 | 3mo > 7mo | 4A |
|  |  | Sex | F(1,16)=7.05 | 0.017 | Males > Females |  |
|  |  | Age x Sex | F(2,32)=0.15 | 0.859 |  |  |
|  | Passive Social | Age | F(2,32)=0.59 | 0.561 |  | 4B |
|  |  | Sex | F(1,16)=1.06 | 0.319 |  |  |
|  |  | Age x Sex | F(2,32)=1.48 | 0.243 |  |  |
|  | Nonsocial Contact | Age | F(2,32)=15.13 | <0.001 | 7mo & 11mo > 3mo | 4C |
|  |  | Sex | F(1,16)=0.53 | 0.476 |  |  |
|  |  | Age x Sex | F(2,32)=12.05 | <0.001 | All ages different in Males only |  |
|  | Nonsocial Attention | Age | F(2,32)=6.80 | 0.003 | 3mo & 7mo > 11mo | 4D |
|  |  | Sex | F(1,16)=10.51 | 0.005 |  |  |
|  |  | Age x Sex | F(2,32)=4.54 | 0.018 | 3mo & 7mo > 11mo in Males only |  |
| Effect of Adolescent Isolation in Males | Active Social | Age | F(2,46)=2.857 | 0.068 |  | 9A |
|  |  | Manipulation | F(1,46)=0.01 | 0.912 |  |  |
|  |  | Age x Manipulation | F(2,46)=0.27 | 0.768 |  |  |
|  | Passive Social | Age | F(2,30)=3.15 | 0.058 |  | 9B |
|  |  | Manipulation | F(1,16)=1.08 | 0.314 |  |  |
|  |  | Age x Manipulation | F(2,30)=0.01 | 0.990 |  |  |
|  | Nonsocial Contact | Age | F(2,30)=32.91 | <0.001 | 11mo > 3mo & 7mo | 9C |
|  |  | Manipulation | F(1,16)=0.18 | 0.676 |  |  |
|  |  | Age x Manipulation | F(2,30)=3.42 | 0.046 | All ages different in Ctrl, but only 11mo > 3mo & 7mo in P31-37iso |  |
|  | Nonsocial Attention | Age | F(2,46)=17.16 | <0.01 | 3mo & 7mo > 11mo | 9D |
|  |  | Manipulation | F(1,46)=0.05 | 0.819 |  |  |
|  |  | Age x Manipulation | F(2,46)=1.50 | 0.233 |  |  |
| Effect of Adolescent Isolation in Females | Active Social | Age | F(2,24)=1.20 | 0.318 |  | 9E |
|  |  | Manipulation | F(1,14)=0.62 | 0.444 |  |  |
|  |  | Age x Manipulation | F(2,24)=1.12 | 0.344 |  |  |
|  | Passive Social | Age | F(2,24)=0.26 | 0.772 |  | 9F |
|  |  | Manipulation | F(1,14)=9.41 | 0.008 | Ctrl > P23-29iso |  |
|  |  | Age x Manipulation | F(2,24)=1.06 | 0.361 |  |  |
|  | Nonsocial Contact | Age | F(2,24)=0.17 | 0.847 |  | 9G |
|  |  | Manipulation | F(1,14)=0.00 | 0.953 |  |  |
|  |  | Age x Manipulation | F(2,24)=0.19 | 0.825 |  |  |
|  | Nonsocial Attention | Age | F(2,24)=0.21 | 0.810 |  | 9H |
|  |  | Manipulation | F(1,14)=0.03 | 0.874 |  |  |
|  |  | Age x Manipulation | F(2,24)=0.29 | 0.753 |  |  |

**Table 5:** Statistical details for Open Field. n.s. = not significant; did not pass statistical significance upon multiple comparisons corrected post-hoc testing

|  | Measurement | ANOVA Factor | Statistic | p-value | Post-hoc | Figure |
| --- | --- | --- | --- | --- | --- | --- |
| Sex Differences in Natural Aging | OF Inner Exploration | Age | $F(1.51, 24.20)=4.57$ | 0.029 | n.s. | 5 |
| | | Sex | $F(1, 16)=51.02$ | <0.001 | Male > Female | |
| | | Age x Sex | $F(2, 32)=0.54$ | 0.589 | | |
| Effect of Adolescent Isolation in Males | OF Inner Exploration | Age | $F(1.50, 22.48)=8.55$ | 0.004 | 3mo < 7mo & 11mo | 10A |
| | | Manipulation | $F(1, 16)=2.04$ | 0.173 | | |
| | | Age x Manipulation | $F(2, 30)=0.27$ | 0.768 | | |
| Effect of Adolescent Isolation in Females | OF Inner Exploration | Age | $F(1.93, 23.10)=6.16$ | 0.008 | 3mo < 11mo | 10B |
| | | Manipulation | $F(1, 14)=0.00$ | 0.956 | | |
| | | Age x Manipulation | $F(2, 24)=1.22$ | 0.314 | | |

### Sex Differences in Natural Social Aging

*Naturally aging male and female rats follow the same social aging trajectory in a Social vs. Object choice test* The Social vs. Object choice test is a simple and widely used assessment of general sociability in rodents [31]. Typically, rodents prefer social over nonsocial stimuli, and thus will spend more time exploring a novel social stimulus vs. a novel object stimulus. We assessed Social vs. Object choice in naturally aging rats with a new sex-matched young adult social stimulus at 3mo, 7mo, and 11mo of age (**Fig. 1**; **Table 1**). In both sexes, time spent in social exploration reduced from 3 to 7mo, while time spent in object exploration increased from 7 to 11mo. There was no effect of sex in social exploration, but females exhibited more overall object exploration than males. The percentage of time the rat spends exploring the social stimulus while in the social half of the arena decreased with age and was higher in females than in males. Interestingly, the percentage of time the rat spends exploring the object stimulus while in the nonsocial half of the arena decreased from 3 to 7mo before returning to 3mo levels at 11mo. This pattern was observed in both sexes.

**Fig. 1:**
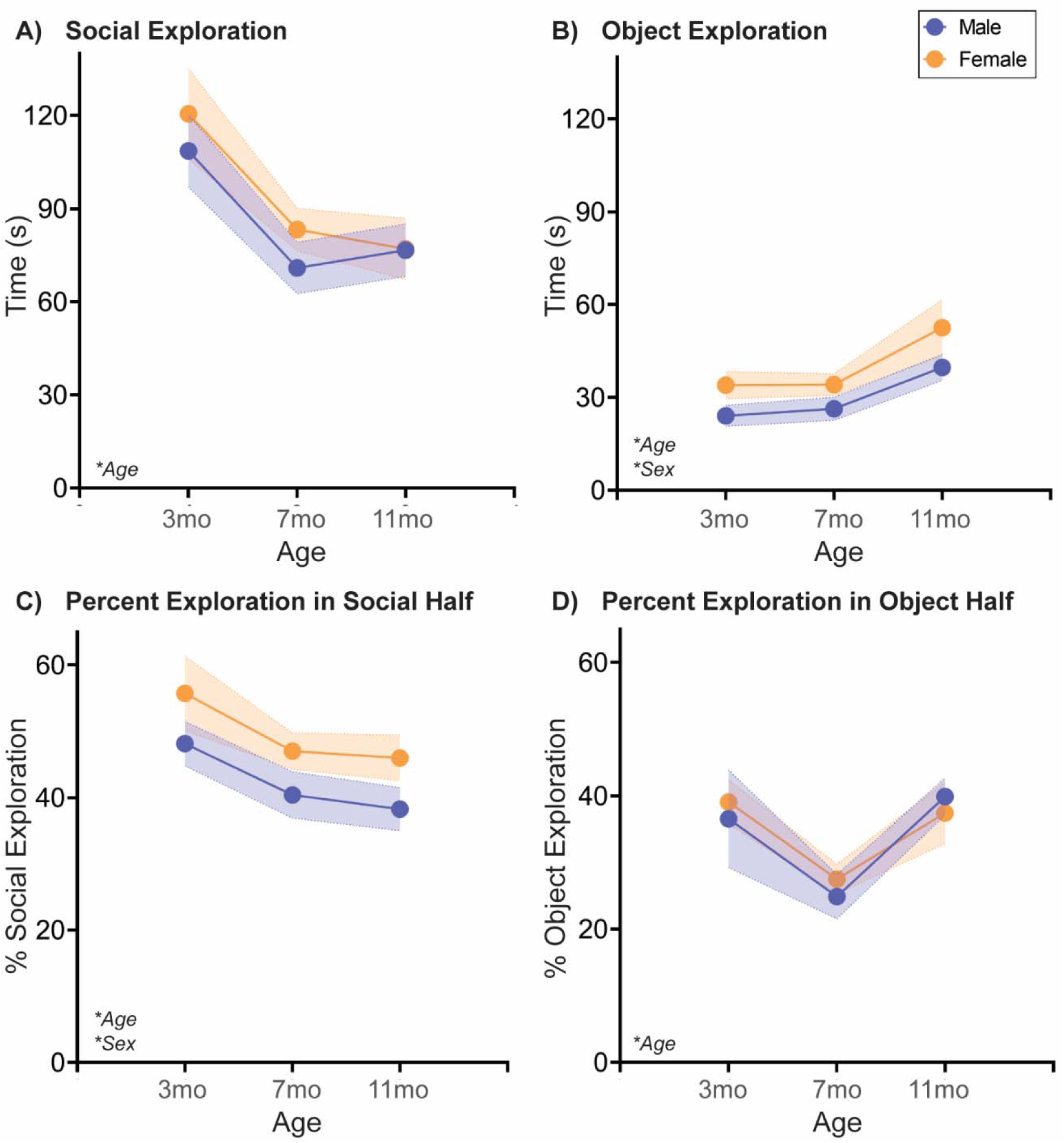
Social exploration decreases, and object exploration increases with age in both sexes in a Social vs. Object choice test. At three different ages in adulthood, rats are placed in a two-chamber compartment with free access to explore a novel adult sex-matched conspecific or a novel nonsocial object for 5 mins. In both sexes, **(A)** social exploration decreased with age, most significantly from 3 to 7 mos, and **(B)** object exploration increased with age, most significantly from 7 to 11 mos. In the latter, females explored the object stimulus more overall than males. **(C)** Both sexes decreased the percentage of time spent in social exploration while in the social half of the arena from 3 to 11 mos, while females had overall higher percentages of social exploration in the social half relative to males. In contrast, **(D)** the percentage of time spent in object exploration while in the object half of the arena decreased from 3 to 7 mos before rebounding at 11 mos. This trend was observed in both sexes. *n*=6-10/sex; test was repeated in the same rats at each time point. Histograms depict average ± SEM. Italicized bottom left text indicates statistically significant (\**p*<0.05) main effects and interactions. All statistical details can be found in **Table 1**.

#### Naturally aging male and female rats follow different social aging trajectories in a Social Novelty Preference choice test

The Social Novelty Preference test is a more complex social choice test in which a rodent is given the option of exploring either a novel, sex-matched young adult social stimulus or a familiar cage mate [32]. Typically, rodents prefer novelty over familiarity, and thus will spend more time exploring a novel social stimulus vs. a cage mate. In this test (**Fig. 2**; **Table 2**), there were statistically significant sex x age interactions in both novel and familiar stimulus exploration, but each sex was responsible for just one effect. In males, but not females, novel social exploration decreased with age (3mo > 11mo). Conversely, in females, but not males, there was a sharp decline in familiar social exploration from 3mo to 7mo that then continued to more modestly decline at 11mo. The percentage of time the rat spends exploring the novel social stimulus while in the novel half of the arena decreased with age in both sexes, with females exhibiting higher percent exploration than males. Aging also decreased percentage of time spent exploring the familiar social stimulus while in the familiar half of the arena, though there was no sex difference in this trajectory.

**Fig. 2:**
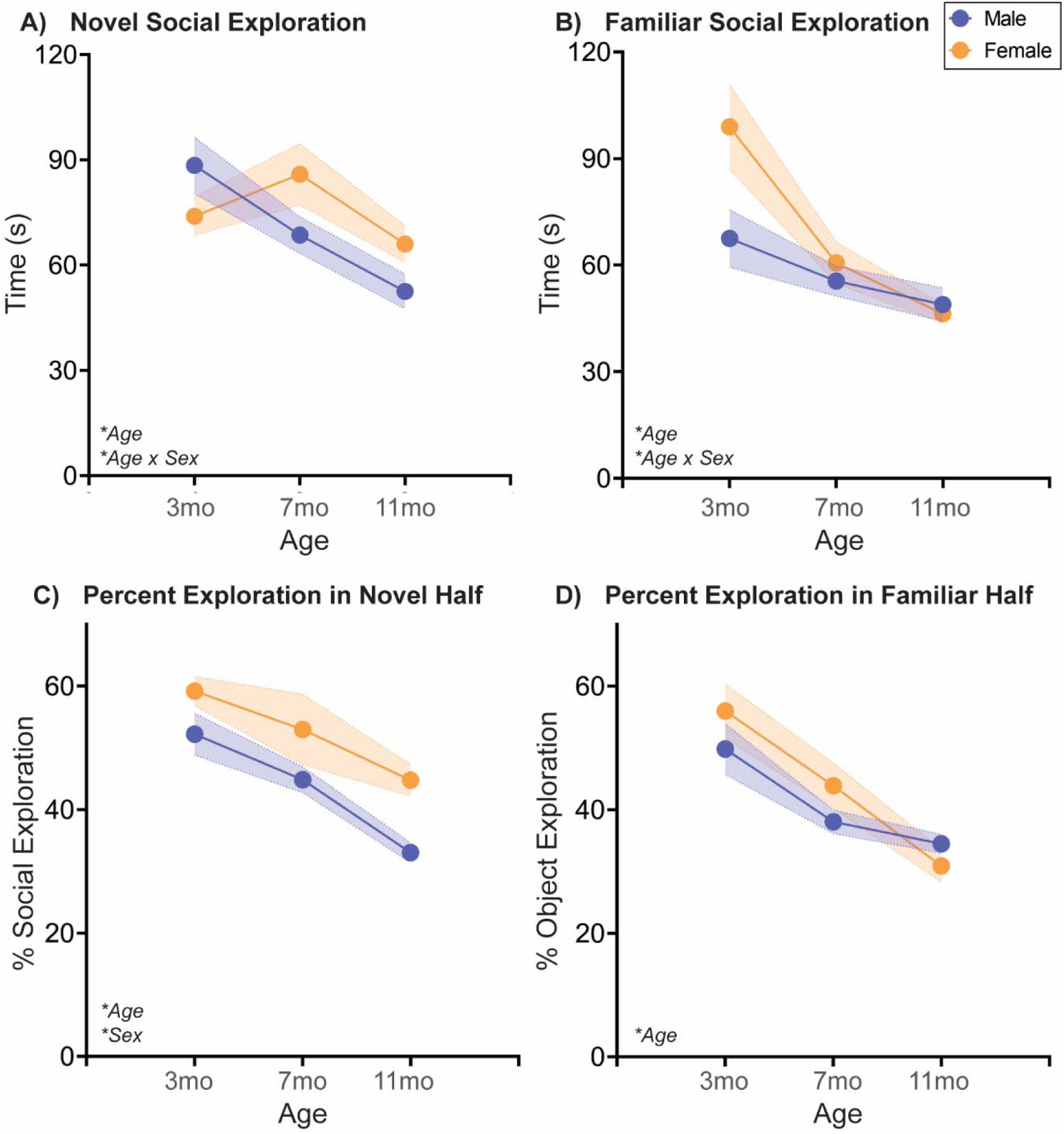
Novel and familiar social exploration decreases with age in sex-specific ways in a Social Novelty Preference choice test. At three different ages in adulthood, rats are placed in a two-chamber compartment with free access to explore a novel adult sex-matched conspecific or a familiar sex- and age-matched cage mate for 5 mins. **(A)** In males, but not females, novel social exploration decreased linearly with age. **(B)** In both sexes, familiar social exploration decreased with age, but by a greater degree from 3 to 7mo in females due to a higher 3mo exploratory baseline. **(C)** Both sexes decreased the percentage of time spent in novel social exploration while in the novel social half of the arena from 3 to 11 mos, while females had overall higher percentages of novel social exploration in the social half relative to males. Similarly, **(D)** both sexes decreased the percentage of time spent in familiar social exploration while in the familiar social half of the arena from 3 to 7 mos. *n*=6-10/sex; test was repeated in the same rats at each time point. Histograms depict average ± SEM. Italicized bottom left text indicates statistically significant (\**p*<0.05) main effects and interactions. All statistical details can be found in **Table 2**.

#### Naturally aging male and female rats follow different social aging trajectories in a Novel Juvenile Interaction test

Assessment of free social interaction with a same-sex pre-pubescent juvenile is common in studies using aged and/or sick rodents [33, 34]. Time spent in interaction with the juvenile typically decreases in sick rodents compared to healthy rodents, which is exacerbated by aging [33]. In our studies (**Fig. 3**; **Table 3**), we measured not just active social interaction, but also other facets of social behavior (described in the Methods; see also **Supp. Fig. 1**). In both male and female rats, both active and passive social interaction declined with age. Nonsocial contact increased with age in males, but not females, while nonsocial attention did not vary significantly by sex or age.

**Fig. 3:**
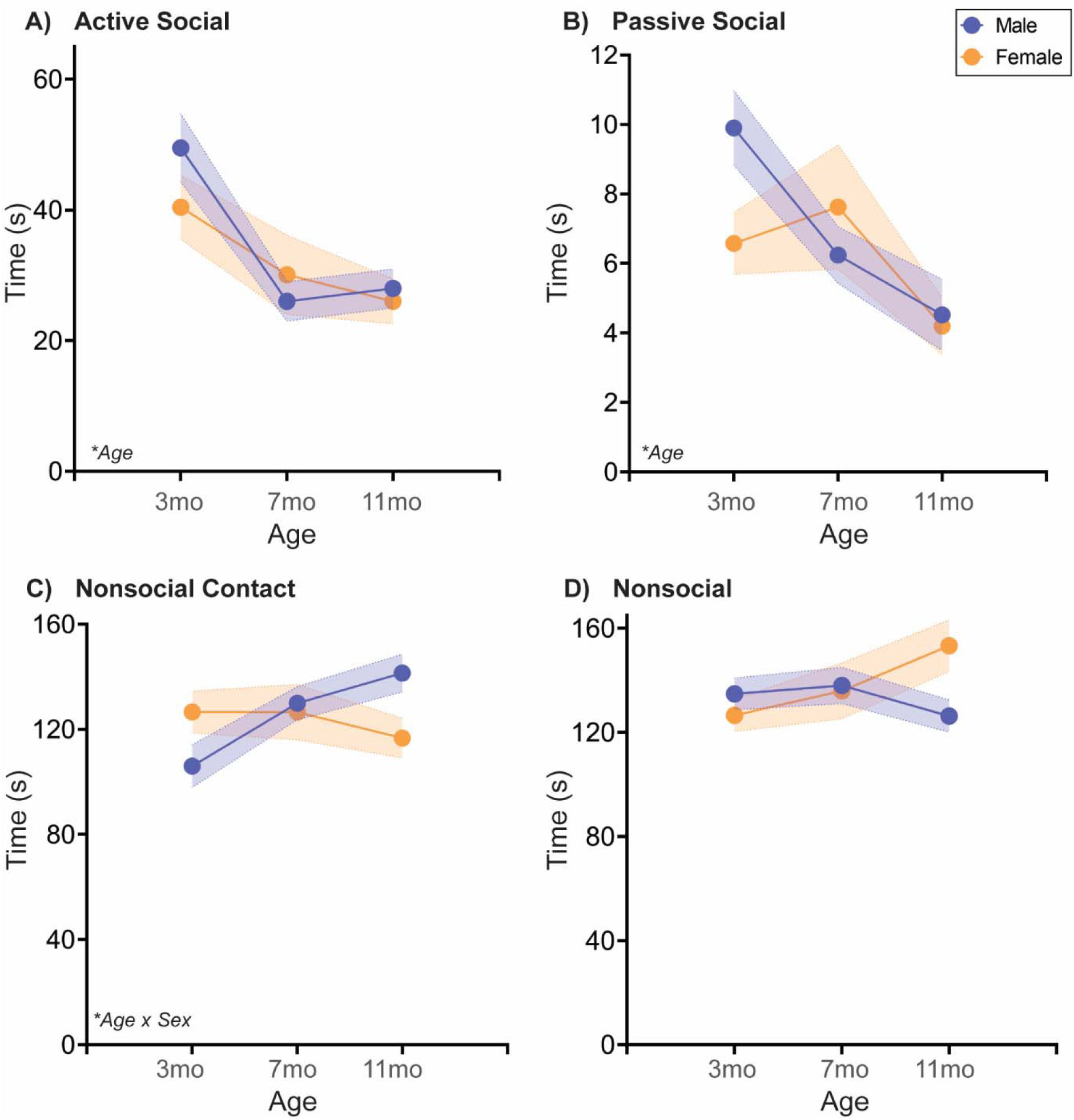
Free social behavior towards a novel juvenile changes sex-specifically with age. At three different ages in adulthood, rats are placed in a clean cage with a novel sex-matched juvenile rat (∼P25-26) and recorded for 5 mins. Behavior of the experimental rat was coded as Active, Passive, Nonsocial Contact, or Nonsocial Attention as depicted in **Supp. Fig. 1**. **(A-B)** Active and passive social behaviors toward the juvenile decreased with age in both sexes. **(C)** In males, but not females, nonsocial contact increased with age, most significantly from 3 to 7 mo, while **(D)** nonsocial attention did not significantly change with age in either sex. *n*=6-10/sex; test was repeated in the same rats at each time point. Histograms depict average ± SEM. Italicized bottom left text indicates statistically significant (\**p*<0.05) main effects and interactions. All statistical details can be found in **Table 3**.

*Naturally aging male and female rats follow different social aging trajectories in a Familiar Interaction test* Finally, one infrequently assessed social relationship is the established familiar relationship within the home cage. In our studies (**Fig. 4**; **Table 4**), familiar cage mates were also siblings and had been pair-housed since weaning. In both male and female rats, active social interaction declined from 3 to 7mo, with males exhibiting higher active social interaction than females. Passive social interaction did not vary significantly by sex or age. Interestingly, nonsocial contact increased with each successive age in males, whereas female nonsocial contact did not change with age. Further, nonsocial attention declined from 7 to 11mo in males, while nonsocial attention did not change with age in females.

**Fig. 4:**
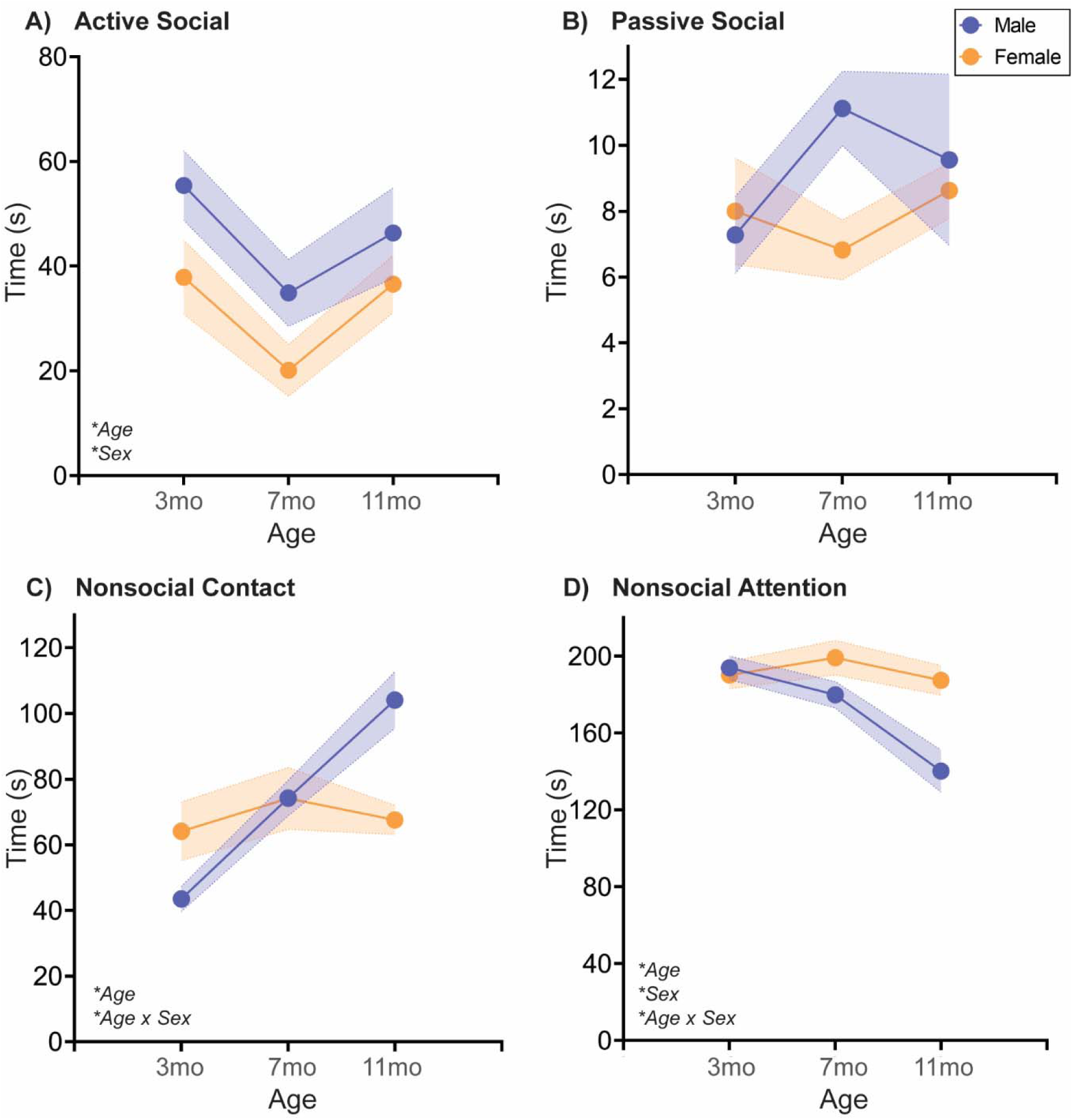
Free social behavior towards a familiar cage mate changes sex-specifically with age. At three different ages in adulthood, age- and sex-matched cage mates are separated for ∼20mins and then placed together in a clean cage and recorded for 5 mins. Behavior of each rat was coded as Active, Passive, Nonsocial Contact, or Nonsocial Attention as depicted in **Supp. Fig. 1**. **(A)** Active social behavior toward a familiar cage mate decreased from 3 to 7mo before rebounding at 11mo. Although both sexes exhibited this pattern, males had overall more active social interaction with their cage mates than females. **(B)** There was no significant regulation of Passive social interaction with age or by sex. In males, but not females, **(C)** nonsocial contact increased linearly with age, while **(D)** nonsocial attention decreased with age, most significantly between 7 and 11mo. *n*=6-10/sex; test was repeated in the same rats at each time point. Histograms depict average ± SEM. Italicized bottom left text indicates statistically significant (\**p*<0.05) main effects and interactions. All statistical details can be found in **Table 4**.

#### Naturally aging male and female rats follow the same aging trajectories in an Open Field test

As a measure of changes in anxiety-like or exploratory behavior, we additionally performed an Open Field test in all rats at each age (**Fig. 5**; **Table 5**). Time spent in the inner zone of the arena, indicative of reduced anxiety-like behavior and/or increased exploratory behavior, increased with age in both sexes, with males spending more time in the inner zone in general.

**Fig. 5:**
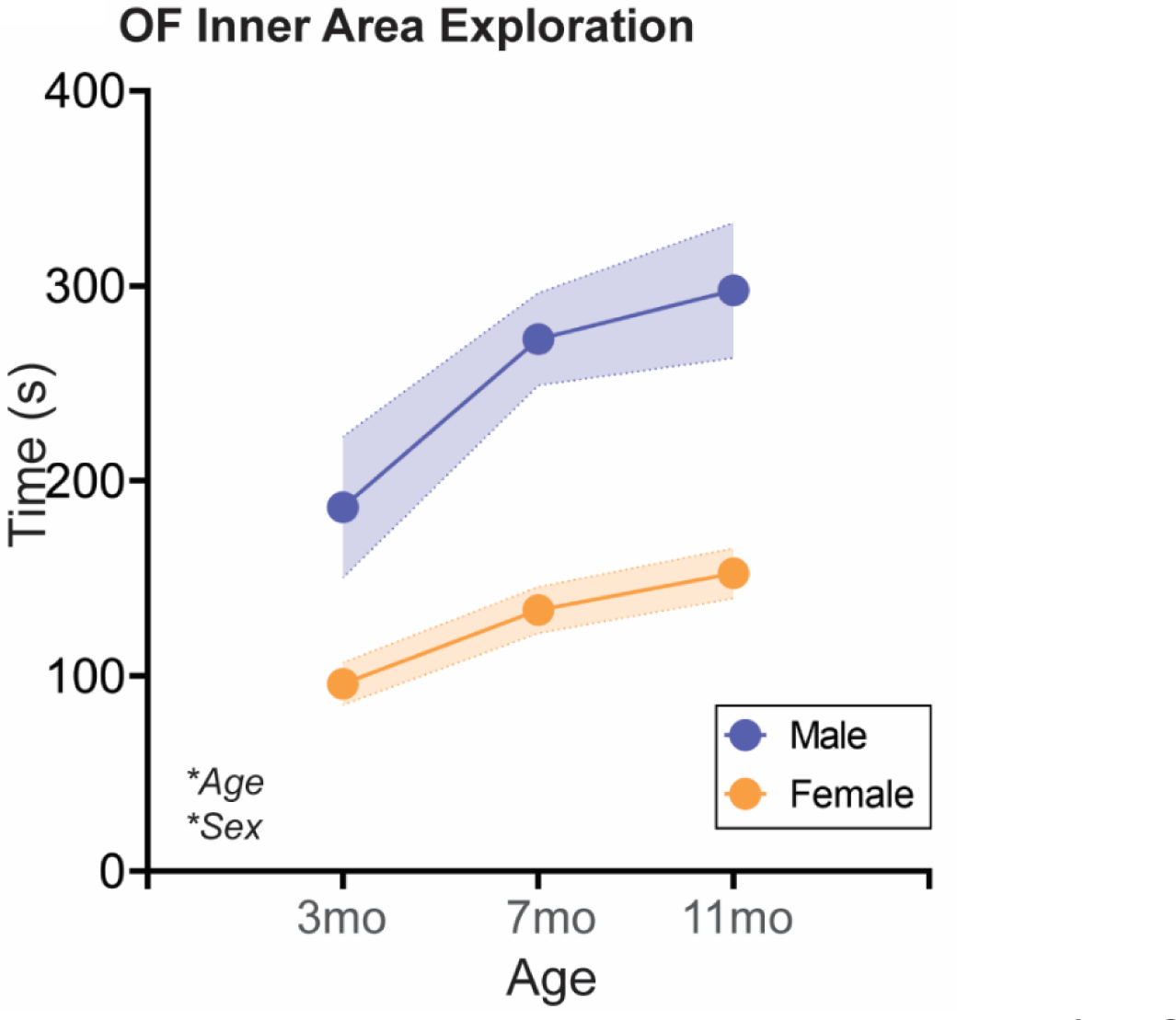
Aging increases time spent in the inner zone of an Open Field apparatus in both sexes. At three different ages in adulthood, rats are placed in a square open field arena with free access to explore for 5 mins. Time spent in the inner zone of the Open Field, associated with reduced anxiety-like behavior and/or increased exploratory behavior, increased with age in both sexes. Males spend more time in the inner zone overall compared to females. *n*=6-10/sex; test was repeated in the same rats at each time point. Histograms depict average ± SEM. Italicized bottom left text indicates statistically significant (\**p*<0.05) main effects and interactions. All statistical details can be found in **Table 5**.

### Effects of Isolation Stress During Sex-Specific Critical Periods of Adolescent Social Development on Social Aging

In addition to naturally aging male and female rats, we had two groups of rats that received social isolation for one week during adolescent critical periods for social development known to vary by sex. Females were single-housed from P23-P30 before re-housing with previous cage mates for the rest of the study, while males were single-housed from P31-38. Thus, males and females were analyzed separately in the following sets of data and we discuss differences in behavioral patterns between the sexes rather than a direct comparison between the sexes at any age.

#### Adolescent isolation stress impacts Social vs. Object choice behavior age-independently, but differently in each sex

Adolescent stress did not impact social aging in most measurements in the Social vs Object choice test, but in both sexes, isolation during the respective sex-specific adolescent critical period impacted the percentage of object exploration when in the object half of the arena. This effect was also sex-specific: in males, P31-P38 isolation reduced percent of object exploration, while in females P23-30 isolation increased percent of object exploration. There were no age x manipulation interactions in the selected measurements, indicating adolescent stress did not impact the social aging trajectory in this test (**Fig. 6**; **Table 1**).

**Fig. 6:**
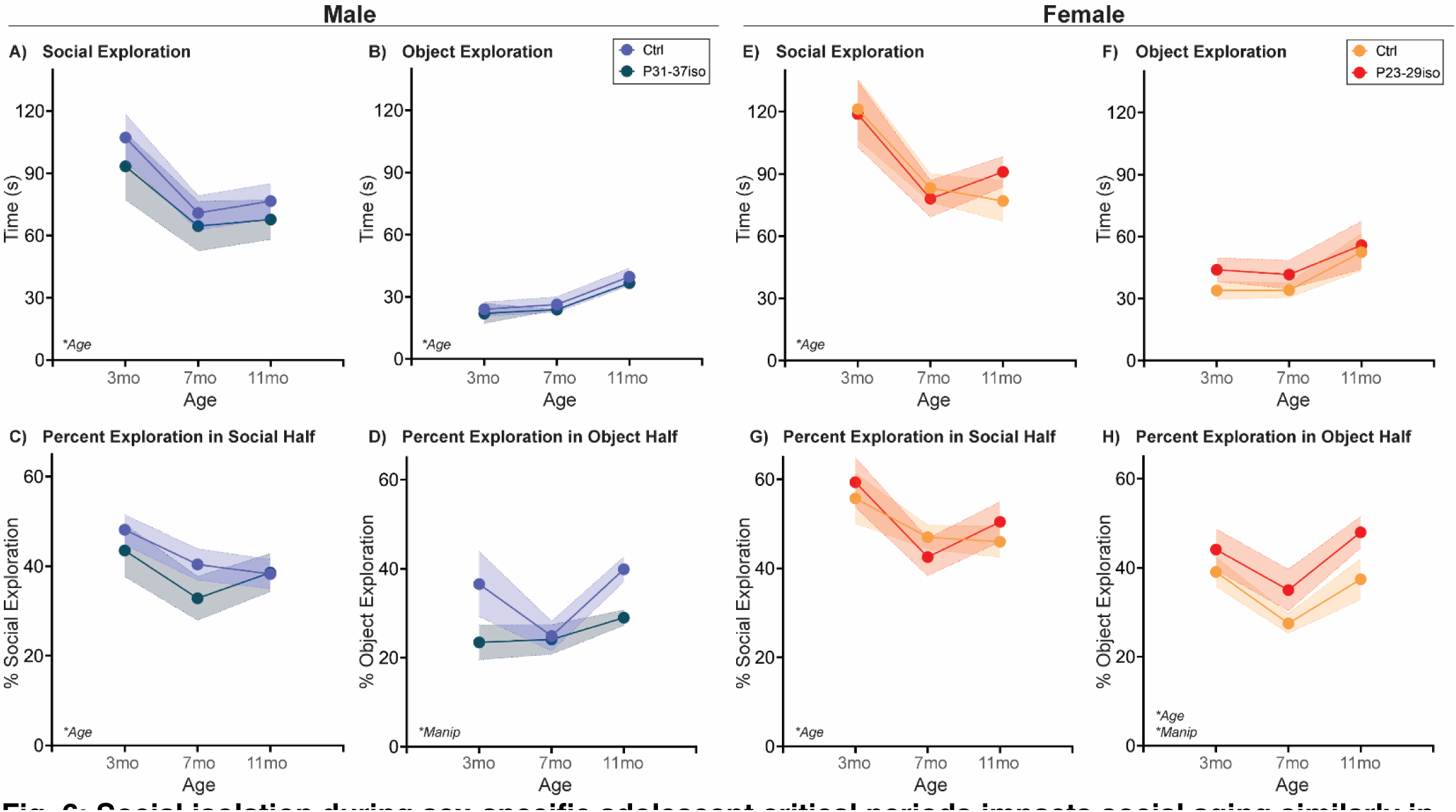
Social isolation during sex-specific adolescent critical periods impacts social aging similarly in both sexes in a Social vs. Object choice test. Rats are single-housed during sex-specific critical periods for adolescent social development, P31-37 in males and P23-29 in females, and then re-housed with previous cage mates. At three different ages in adulthood, rats are placed in a two-chamber compartment with free access to explore a novel adult sex-matched conspecific or a novel nonsocial object for 5 mins. **(A-B, E-F)** In both sexes, social exploration decreased with age and object exploration increased with age. The latter did not reach statistically significance in females, though the same behavior pattern is present. Adolescent isolation did not impact exploration of either stimulus. **(C, G)** In both sexes, aging decreased the percentage of time spent in social exploration while in the social half of the arena from 3 to 11 mos, with no effect of adolescent isolation. **(D, H)** In both sexes, adolescent isolation did significantly regulate the percentage of time spent in object exploration while in the object half of the arena. However, in males, adolescent isolation reduces the overall percent of object exploration while in the object half, whereas in females adolescent isolation increases the overall percent of object exploration while in the object half. *n*=6-10/sex; test was repeated in the same rats at each time point. Histograms depict average ± SEM. Italicized bottom left text indicates statistically significant (\**p*<0.05) main effects and interactions. “Manip” = adolescent manipulation. All statistical details can be found in **Table 1**.

#### Adolescent isolation stress impacts the social aging trajectory in the Social Novelty Preference choice test in females, but not males

In males, there was a decrease in all measurements with age, with no significant effects of adolescent social isolation during the male critical period for social development. In females, there was a decrease in all measurements with age, but an additional age x manipulation interaction in time spent exploring the familiar social stimulus. In naturally aging females, familiar social exploration decreases with age, while in females subjected to social isolation during their adolescent critical period, there was no statistically significant change in familiar stimulus exploration with age (**Fig. 7**; **Table 2**).

**Fig. 7:**
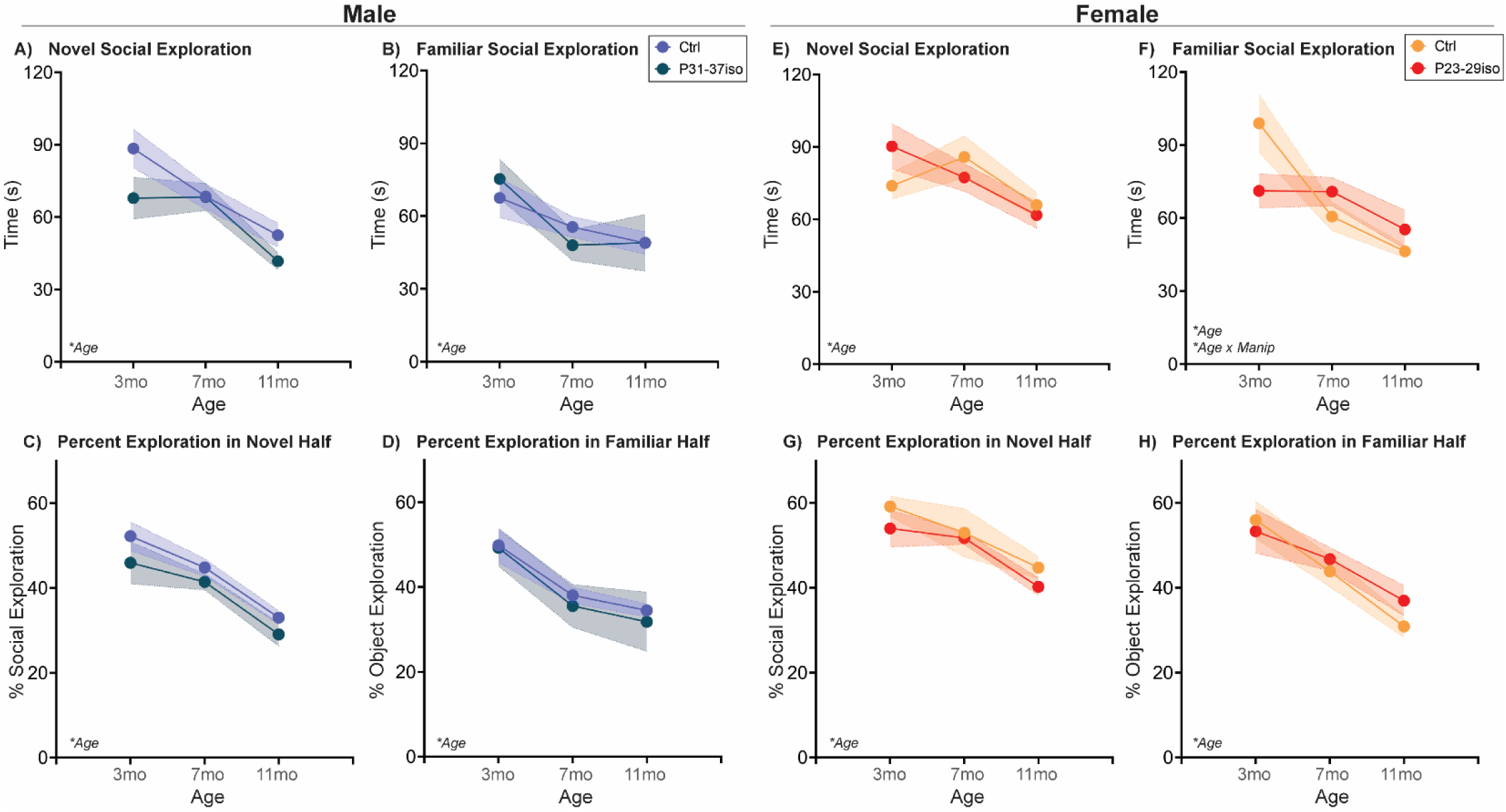
Social isolation during sex-specific adolescent critical periods impacts social aging in females, but not males, in a Social Novelty Preference test. Rats are single-housed during sex-specific critical periods for adolescent social development, P31-37 in males and P23-29 in females, and then re-housed with previous cage mates. At three different ages in adulthood, rats are placed in a two-chamber compartment with free access to explore a novel adult sex-matched conspecific or a familiar age- and sex-matched cage mate for 5 mins. **(A-D)** In males, exploration of both stimuli and percent exploration while in each respective compartment decreased with age and was not impacted by adolescent isolation. **(E, G-H)** Similarly, in females, exploration of the novel social stimulus and percent exploration of the novel and familiar social stimuli while in each respective compartment decreased with age independent of adolescent isolation. **(F)** However, in females only there was an interaction between age and manipulation with respect to familiar social exploration such that adolescent isolation induced lower 3mo exploration than control, and consequently no decline in exploration from 3 to 7mo that is observed in control females. *n*=6-10/sex; test was repeated in the same rats at each time point. Histograms depict average ± SEM. Italicized bottom left text indicates statistically significant (\**p*<0.05) main effects and interactions. “Manip” = adolescent manipulation. All statistical details can be found in **Table 2**.

#### Adolescent isolation stress does not impact Novel Juvenile Interaction behavior in either sex

In male rats, aging reduced active and passive social interaction while increasing nonsocial contact. In females, age reduced active social interaction. There were no effects of adolescent isolation for either sex (**Fig. 8**; **Table 3**).

**Fig. 8:**
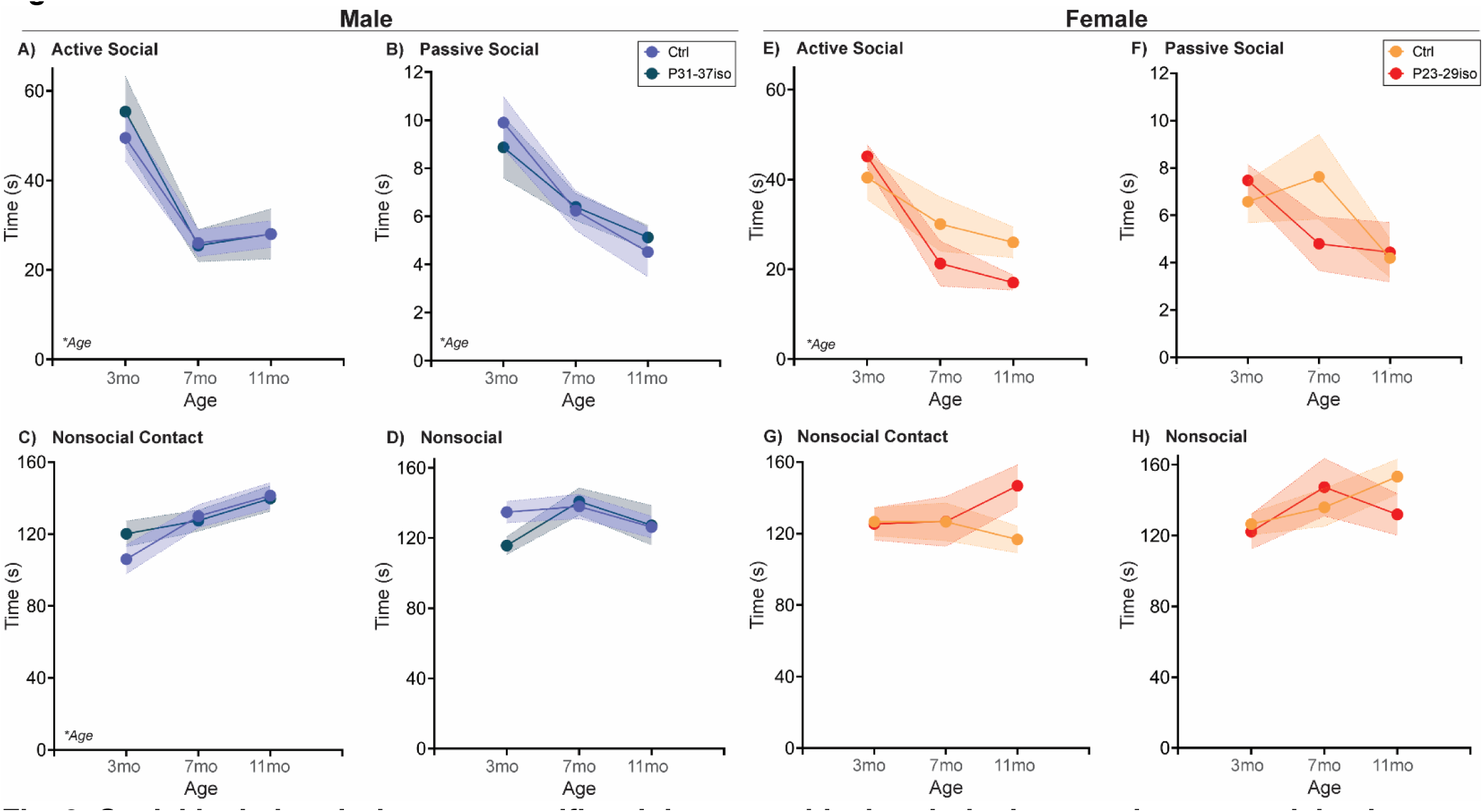
Social isolation during sex-specific adolescent critical periods does not impact social aging behavior towards a novel juvenile. Rats are single-housed during sex-specific critical periods for adolescent social development, P31-37 in males and P23-29 in females, and then re-housed with previous cage mates. At three different ages in adulthood, rats are placed in a clean cage with a novel sex-matched juvenile rat (∼P25-26) and recorded for 5 mins. Behavior of the experimental rat was coded as Active, Passive, Nonsocial Contact, or Nonsocial Attention as depicted in **Supp. Fig. 1**. **(A-D)** In males, Active and Passive social behaviors towards the juvenile decreased with age, Nonsocial contact increased with age, and Nonsocial behavior did not significantly change with age. Adolescent isolation did not impact any of these behavioral measurements. **(E-H)** In females, Active social interaction towards the juvenile decreased with age, while Passive interaction, Nonsocial Contact, and Nonsocial Attention did not change with age. Similar to males, adolescent isolation did not impact any of these behavioral measurements. *n*=6-10/sex; test was repeated in the same rats at each time point. Histograms depict average ± SEM. Italicized bottom left text indicates statistically significant (\**p*<0.05) main effects and interactions. All statistical details can be found in **Table 3**.

#### Adolescent isolation stress impacts Familiar Interaction behavior age-dependently in males, and age-independently in females

In male rats, passive social interaction increased with age and nonsocial attention decreased with age, with no significant effects of adolescent social isolation during the male critical period for social development. However, there was an age x manipulation interaction in nonsocial contact. In naturally aging males, nonsocial contact increased successively with each age, while in males subjected to social isolation during their adolescent critical period, there was increased nonsocial contact only between 7 and 11 mos. In females, there was no impact of age in any measurement, but females subjected to social isolation during their adolescent critical period had reduced passive social interaction age-independently (**Fig. 9**; **Table 4**).

**Fig. 9:**
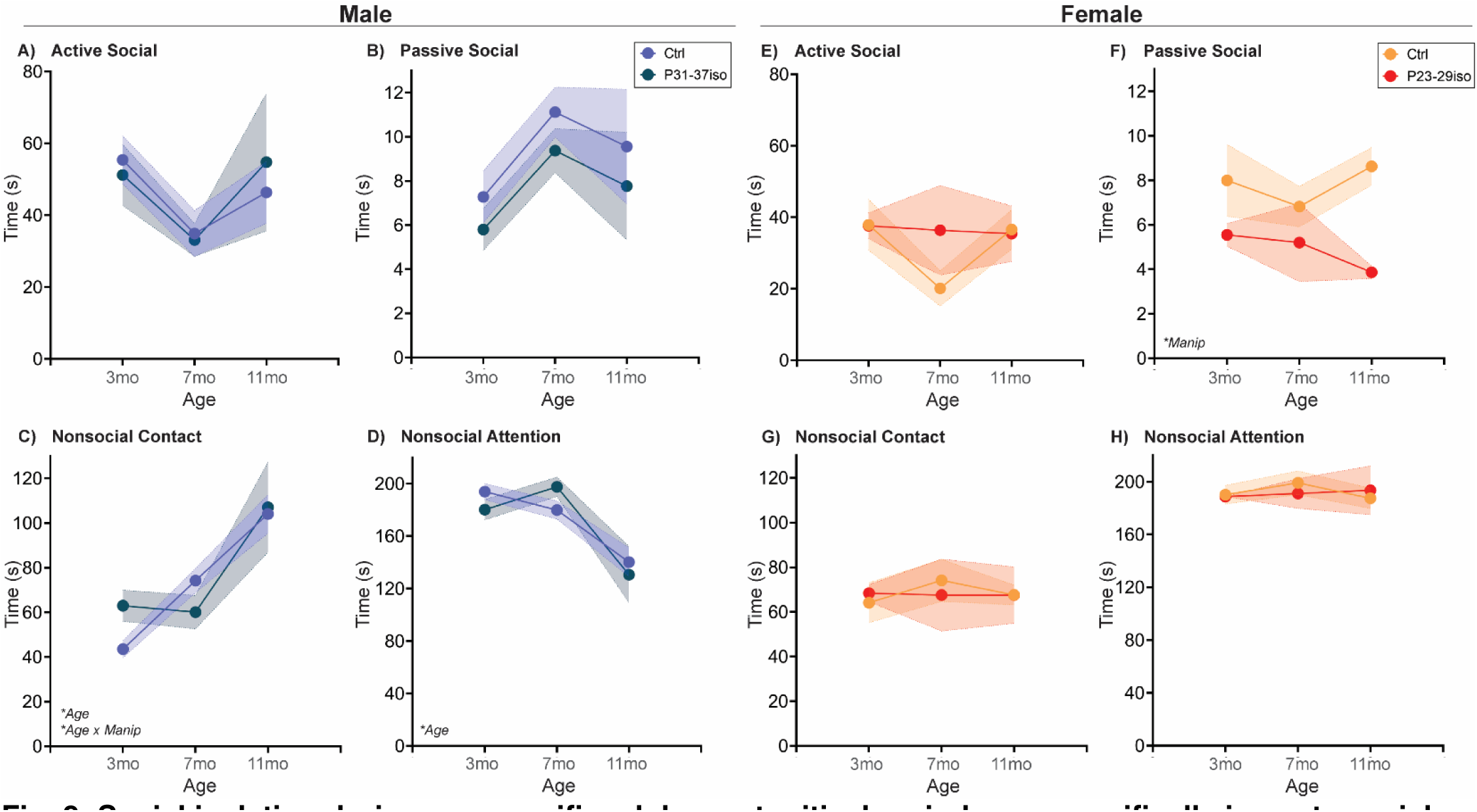
Social isolation during sex-specific adolescent critical periods sex-specifically impacts social aging behavior towards a familiar cage mate. Rats are single-housed during sex-specific critical periods for adolescent social development, P31-37 in males and P23-29 in females, and then re-housed with previous cage mates. At three different ages in adulthood, age- and sex-matched cage mates are separated for ∼20mins and then placed together in a clean cage and recorded for 5 mins at three different ages. Behavior of the rats was coded as Active, Passive, Nonsocial Contact, or Nonsocial Attention as depicted in **Supp. Fig. 1**. **(A-B)** In males, Active and Passive social behaviors with familiar cage mates did not significantly change with age. **(C)** In male Nonsocial Contact behavior there was a age by manipulation interaction such that rats that experienced adolescent isolation had higher Nonsocial Contact than controls at 3mo and thus did not significantly increase this behavior from 3 to 7mo as in control groups. **(D)** In males, Nonsocial Attention decreased with age independent of adolescent isolation. **(E, G-H)** In females, there was no significant regulation of Active social interaction, Nonsocial Contact, or Nonsocial Attention by age or adolescent isolation. **(F)** However, adolescent isolation significantly reduced the overall level of Passive social interaction relative to control females. *n*=6-10/sex; test was repeated in the same rats at each time point. Histograms depict average ± SEM. Italicized bottom left text indicates statistically significant (\**p*<0.05) main effects and interactions. “Manip” = adolescent manipulation. All statistical details can be found in **Table 4**.

#### Adolescent isolation stress does not impact Open Field behavior in either sex

In both sexes, aging increased time spent in the inner zone of the arena, which was not significantly regulated by adolescent isolation in either sex (**Fig. 10**; **Table 5**).

**Fig. 10:**
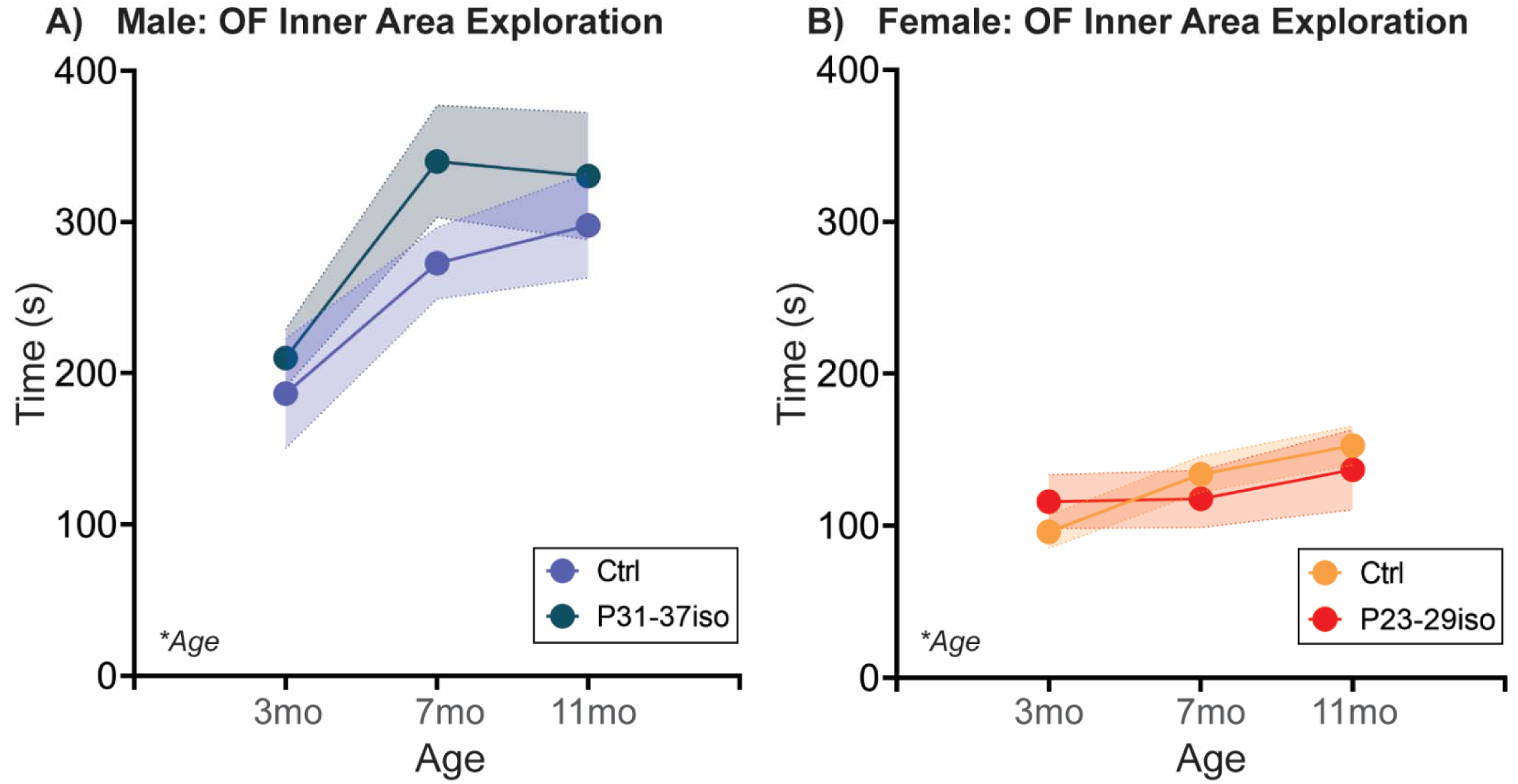
Social isolation during sex-specific adolescent critical periods does not impact time spent in the inner zone of an Open Field apparatus. Rats are single-housed during sex-specific critical periods for adolescent social development, P31-37 in males and P23-29 in females, and then re-housed with previous cage mates. At three different ages in adulthood, rats are placed in a square open field arena with free access to explore for 5 mins. Time spent in the inner zone of the Open Field increased with age in **(A)** males and **(B)** females, and adolescent isolation did not impact this behavior in either sex. Histograms depict average ± SEM. Italicized bottom left text indicates statistically significant (\**p*<0.05) main effects and interactions. All statistical details can be found in **Table 5**.

## DISCUSSION

In this set of studies, we determined how sex impacted natural social aging from 3 to 11mo of age in rats, and assessed whether isolation stress during known critical periods of social development during adolescence would alter the social aging trajectory. We observed clear social aging signatures in all tests administered, but sex differences in natural social aging were most robustly observed when a familiar social stimulus was included in the test. We also observed that adolescent isolation did impact the social behavior, in both age-independent and age-dependent ways, that were entirely sex-specific.

### Natural aging decreases social behavior across tasks

As discussed in the Introduction, social behavior declines with aging across species, but there are numerous documented benefits to maintaining social activity. To our knowledge, this is the first study examining social aging in both sexes and in multiple social tests, concurrently. Our data agree with the literature, as we observed social decline with aging in all tests, including decreased social exploration in the Social vs. Object choice test (**Fig. 1**), decreased novel and familiar social exploration in the Social Novelty Preference choice test (**Fig. 2**), and reduced active social interaction in Juvenile Social Interaction (**Fig. 3**) and Familiar Social Interaction tests (**Fig. 4**). These changes are unlikely to be due to a general increase in anxiety-like behavior and/or decrease in exploratory behavior, as the Open Field data suggest anxiety-like behavior decreases and/or exploratory behavior increases (**Fig. 5**). Peripheral inflammation can also change social behavior and is increased during aging, but we observed no differences in peripheral inflammation in serum or cardiac tissue at the termination of the studies (∼13mo; **Supp. Fig. 1, 2**).

However, both free social interaction tests revealed more complex, male-specific behavioral patterns in which nonsocial contact behavior increased (**Fig. 3,4**), and in Familiar Social Interaction this increase was at the expense of nonsocial behavior. We quantified social interaction into four categories that (in our opinion) vary on the social spectrum: active social interaction is the most social behavior, followed by passive social interaction, followed by nonsocial contact, followed finally by true nonsocial behavior. If these arbitrary, and anthropomorphized designations hold true for a rat, then increased nonsocial contact, especially at the expense of nonsocial attention, would be indicative of increased sociability with aging, in males only.

Regardless, these data would implicate familiar social touch in a male-specific aging phenotype. Social touch is becoming more widely recognized as a powerful pain and stress reliever, with recent data indicating that social touch activates the oxytocin and dopaminergic neural systems to promote its effects [35–37]. Aging is often accompanied by increased pain, which is reported to be higher in women [38]. Could this sex difference be due to male rodents ‘self-medicating’ with social touch [39]?

### Social tests that include familiar conspecifics reveal that males and females exhibit different social aging trajectories

Males and females express social behaviors differently, and also value social behaviors differently, both from an ethological and sociocultural perspective [19–21]. It would thus be unsurprising if males and females exhibited different social aging trajectories. Interestingly, sex differences in social aging were highly task-specific. There were no sex x age interactions in Social vs. Object choice tests, one of the most widely used social tests in the literature. In another common social test specifically in aged rodents, the Juvenile social interaction test, there was a sex x age interaction in which males, but not females: increased nonsocial contact with their juvenile partners with age. However, qualitative examination of the videos suggested the experimental rats of both sexes were eager to be rid of their juvenile counterparts. Let’s be honest, little kids are known to be annoying from time to time. I am allowed to say this because I have a 2.5 year old – don’t @ me. The juvenile rats were crawling over and under the adults and despite repeated, forceful ‘signals’ (read: kicks in the head), would not leave well enough alone. One might predict that female rats might have an ethological drive to be more tolerant with the juveniles, but this did not appear to be the case. Qualitatively, the adult females generated a lot more head kicks than the adult males. We included this test because of its use in the aging literature, but we would thus argue this is a stressed social interaction *at best*. It will be interesting to determine whether novel adult conspecifics would elicit more appropriate social behaviors in a free interaction context, and thus be more likely to reveal pertinent social aging changes. FWIW, in our hands, conventional analysis of novel social interaction between sex-matched adult rats is infrequently informative [30].

What is far more informative, as is the case in the present dataset, is when a familiar conspecific is involved. In Social Novelty Preference, exploration of the novel social stimulus declined with age in males, with no significant change in females, while exploration of the familiar social stimulus declined with age in females, with no significant change in males. In the Familiar interaction test, nonsocial contact increased and nonsocial attention decreased with age in males, with no significant regulation in females in either parameter. So what’s different about familiar social stimuli? Generally familiar social stimuli evoke less active social interaction than novel social partners, which may create a larger parameter space in which those behaviors can change. There are also instances in which familiar social stimuli are reported to be preferred, e.g. in the context of sickness [40]. There are certainly very different roles that a novel vs. familiar social partner would play in one’s life, but to our knowledge, there is little known about the relative contribution of novel vs. familiar social engagement in the context of aging – or really any neurological disorder. Not even the extensive pair bonding literature in voles would necessarily be relevant in this instance, as those social connections are in breeding pairs. At least from my experience, my siblings are some of the most important and influential people in my life. One interesting study found that housing with a novel mouse upon weaning vs. housing with same-sex siblings persistently negatively impacted biology and behavior into adulthood, in males but not females [41]. At weaning! One would expect that having all of adolescence to form that social bond would make up for the unfamiliarity of the partner. That appears to not be the case. Our data would suggest this might be an important area to study in the future.

### Short-term isolation stress restricted to adolescent critical periods for social development is sufficient to impact age-independent social behavior *and* social aging in sex-specific ways

Adolescent development is quite protracted, and can last from 10-24 years in humans and from P21-P60 in rats and mice [42]. It is a developmental period in which organisms shift from family-centered social interactions to peer-centered social interactions, which is accompanied by increased exploratory behavior and risk-taking as well as increased susceptibility to peer pressure [23, 43, 44]. Social isolation during adolescence has long been known to produce negative consequences, some of which are sex-specific [26]. However, in most of this literature the isolation stress is administered throughout the entire adolescent period, which is neither translationally relevant nor particularly helpful in terms of assigning causality to specific developmental mechanisms that might be altered by adolescent stress. In our model, we limited social isolation to one week in each sex, during periods known to be critical for social development (P23-30 in females, P31-38 in males; we will discuss this further in the forthcoming *Mechanism* section). While this is unlikely to induce the same magnitude and array of behavioral changes as P21-P60 isolation, we would argue this might be a more strategic and informative model for future studies.

We observed age-dependent and age-independent social changes in response to adolescent isolation, all of which were sex-specific. The age-independent main effects of adolescent social isolation were primarily observed in the Social vs. Object choice test. In both sexes, adolescent isolation significantly impacted the percentage of time spent exploring the object stimulus while the rat was in the object half of the arena. In males, this percentage was decreased by adolescent isolation, while in females the percentage was increased. We interpret this measurement to reflect the relative interest evoked by the stimulus. For example, a rat could enter the object half of the compartment simply to explore the arena without paying any attention to the object stimulus. This would result in a (relatively) low percentage of time spent in object exploration while in that half of the arena. However, if the rat was actively interested in the object stimulus, the percent of time spent in exploration while in this half of the arena would be increased. The latter would also be true if there was very little exploration of the arena, and instead the rat made a direct approach to the stimulus. So we think about this measurement as the ‘attractiveness or draw’ of the stimulus. Of course, YMMV and there are a million interpretations that could also be plausible. However, if our interpretation is in the ballpark of appropriate, then adolescent isolation would decrease the attractiveness or draw of the novel object in males, and increase the attractiveness or draw of the novel object in females. One might be able to test such a theory by conducting novel object recognition (NOR) tests. If our working theory is correct, males that experienced P31-38 social isolation might have reduced NOR memory or acquisition exploration, and females that experienced P23-30 social isolation might have increased NOR memory or acquisition exploration. This would be particularly interesting, as there are known deficits in NOR memory that accompany aging in rodents – might this be a stimulus-dependent effect rather than memory decline per se? The relevance of this finding, particularly to social aging, is unclear at this point, but it would be useful to know if this measurement, derived from a fairly simple and widely used test, could be a behavioral biomarker of adolescent stress impacting long-term behavioral outcomes. Because it was age-independently observed, this ‘biomarker’ would be evidence in young adulthood with appropriately powered group sizes. There were no other age-independent effects of adolescent isolation in Social Novelty Preference or Juvenile social interaction, and only a reduction in the already low expression in passive social interaction, in females only, in the Familiar interaction test.

However, there were age x manipulation interactions in the data for both sexes – and again only when familiar social stimuli were included in the tests. The impact on females was evident in Social Novelty Preference: adolescent isolation reduces 3mo familiar exploration similar to control 7mo levels. Thus, the control females decline in this parameter between 3 and 7mo, while the females that were isolated during adolescence were flat between 3 and 7mo. In both groups, there was a further decline from 7 to 11mo. In males performing the Familiar interaction test, adolescent isolation increased nonsocial contact at 3mo similar to 7mo control levels. As control male rats’ nonsocial contact increased from 3 to 7mo, males that received adolescent isolation were unchanged, and then both groups increased nonsocial contact from 7 to 11mo. While the effect of both age x manipulation interactions presented in tests with familiar conspecifics, it may be relevant that each sex displayed altered behavior in a different test, females in Social Novelty Preference and males in Familiar interaction. There are many differences between these tests, but two that stand out are: 1) choice and 2) a shared history. Re: the former, in the Social Novelty Preference test, the option to explore the familiar conspecific, the parameter in which females showed an age x manipulation interaction, is weighed against the option to explore a novel conspecific. It may be the case that females will show more effects of adolescent isolation in situations which there are opposing choices for behavioral allocation. The positive interpretation of each stimulus in a vacuum may not be affected, but the relative positive interpretation is altered. Re: the latter, males performing the Familiar interaction test have the same history of either no manipulation during adolescence or isolation during adolescence. If there are changes in behavior that are too subtle to detect in interaction with typically-developing rats, then placing two subtly impaired rats together may magnify the deficits. Or, as discussed above, perhaps the life-long bond between these two rats just results in an entirely different repertoire of social behaviors. Regardless of the underlying neurobehavioral mechanisms, in both these cases the isolated male and female rats had reached levels of behavior in line with control rats *of an advanced age*. This may suggest that adolescent social isolation is capable of hastening the social aging progression. There are limited data that social stress throughout adolescence reduces lifespan [1] and worsens aging-induced memory decline in mice [13]. We did not observe any lifespan changes by adolescent stress (*data not shown*), but the rats were also not nearing the end of their natural lifespan when we stopped our studies. Because aging is associated with social decline, which is in turn detrimental to mental and physical health, our data might suggest that rats that experienced adolescent isolation may be less mentally or cognitively fit at younger ages, which then becomes occluded by natural aging processes. This may be an important area of study, as there are a lot of social relationships that are established or mature in young adulthood that may play lifelong roles in health, quality of life, and longevity.

### Social Aging is not linear and may begin earlier in life than expected

When we began this study, we considered what ‘social aging’ would look like: would there be a linear decline in sociability? Stable social behaviors until our most advanced aging time point? Would we see anything at all given that we ended the study in ‘advanced middle age’ for a rat? What social aging will look like will depend on the test and sex, but based on our data, there was quite a lot of social change between 3 to 7 mo, and then more behavioral similarity between 7 and 11 mo. This was surprising, because if there was going to be a nonlinear pattern I would have assumed 3 and 7 mo would have been more similar, and both quite different from 11 mo. Furthermore, when adolescent isolation did impact the social phenotype in an age-dependent manner, in both sexes it appeared to accelerate the 3-7mo profile without impacting the 7-11mo change. The rats in these studies were bred and raised in-house, so there were no substantial changes in their daily life between 3 and 7 mo that can (to our knowledge) account for these changes. However, there are data suggesting that social changes associated with aging do occur prior to the more obvious cognitive defects that emerge with aging [45]. It will be interesting in future studies to determine whether a pro-social intervention (whatever that means) between 3 and 7 mo would be particularly beneficial for social aging outcomes. If yes, our data would also predict that the same pro-social outcome would not be as impactful between 7 and 11mo.

### Possible Mechanisms

There are a lot of really important neurodevelopmental changes occurring in many different brain regions during adolescence. However, we favor one mechanistic explanation for the observed effects of adolescent isolation: we previously published that microglia, the resident immune cells of the brain, mediate synaptic pruning in the nucleus accumbens (NAc) reward region during the sex-specific periods we targeted in these studies (P20-30 in females and P30-40 in males) [28]. Inhibiting microglial pruning in the NAc at the beginning of each respective pruning period increased social play in both sexes [28], an adolescent-typical social behavior that declines during maturation into young adulthood. These data suggest that this immune process mediates the natural developmental decline in immature social behaviors. One might predict that a stressor occurring coincidently with this ongoing immune process might be pro-inflammatory, and thus exacerbate pruning, in turn reducing sociability into adulthood. The effects of stress on microglial activity are complex, and depend on the nature of the stressor, the life period the stressor is applied, and the sex of the organism [46]. But this is where I would be looking first if I were to follow-up on these data. We did begin to examine whether there was increased inflammation in the periphery (serum and cardiac tissue), but did not see any changes by sex or by adolescent manipulation (**Supp. Figs. 2-3**).

## Acknowledgements

**Acknowledgements:** This work was supported by the National Institutes of Health R01DA052889 and R03AG07011 to AMK and Albany Medical College Start-up funds to AMK. We thank Dr. Justin Bourgeois for assistance with tissue collection.

**Author Declarations:** AMK designed the experiments. CF, JMK, IP, and DNK performed the experiments. ELE and AMK analyzed the experiments. AMK wrote the manuscript. All authors edited the manuscript. The authors declare no conflicts of interest.

---

**Supp. Fig. 1:**
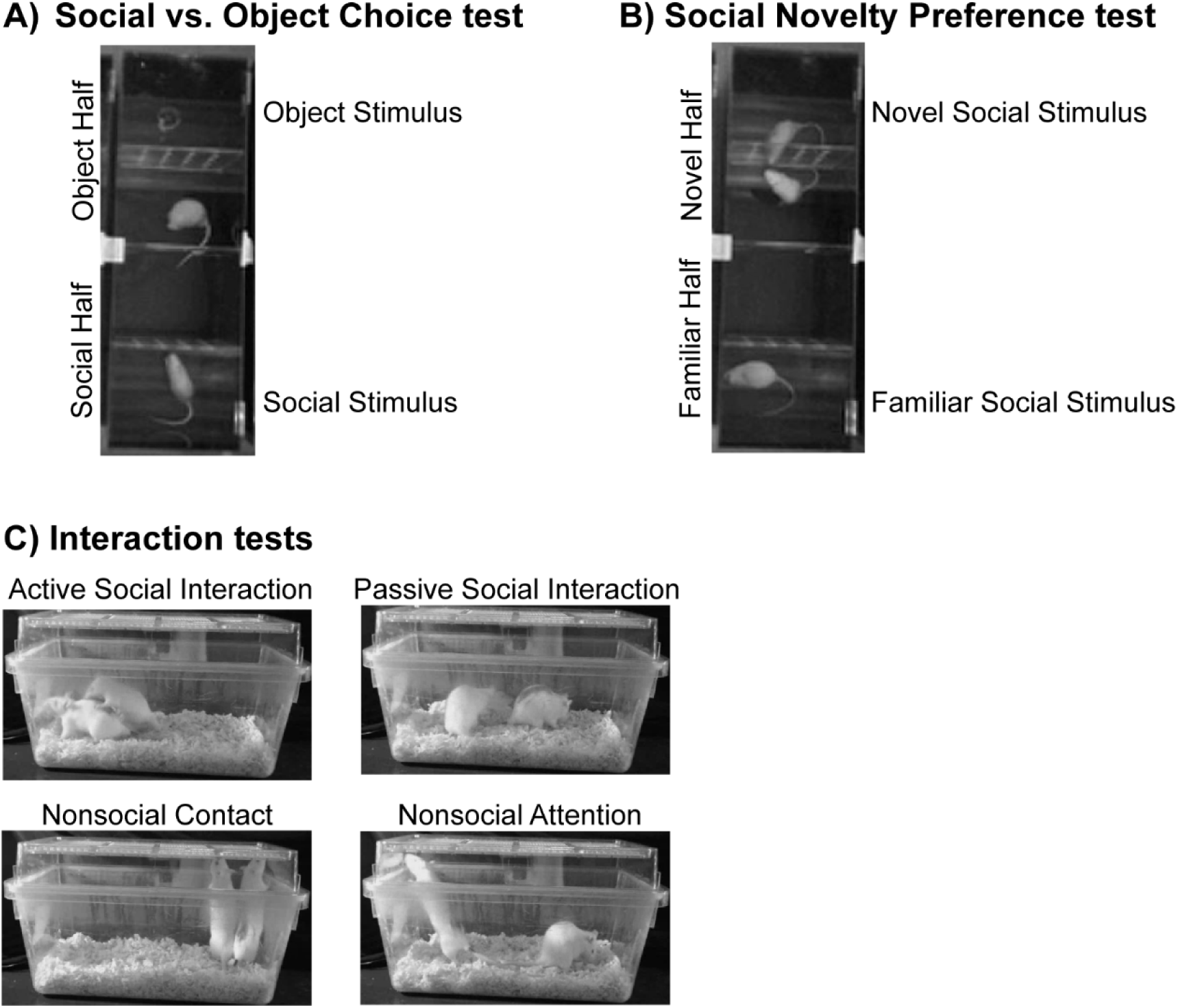
Representative images of social tests. Depiction of social choice tests **(A-B)** and behavioral quantification of Interaction tests **(C)**.

**Supp. Fig. 2:**
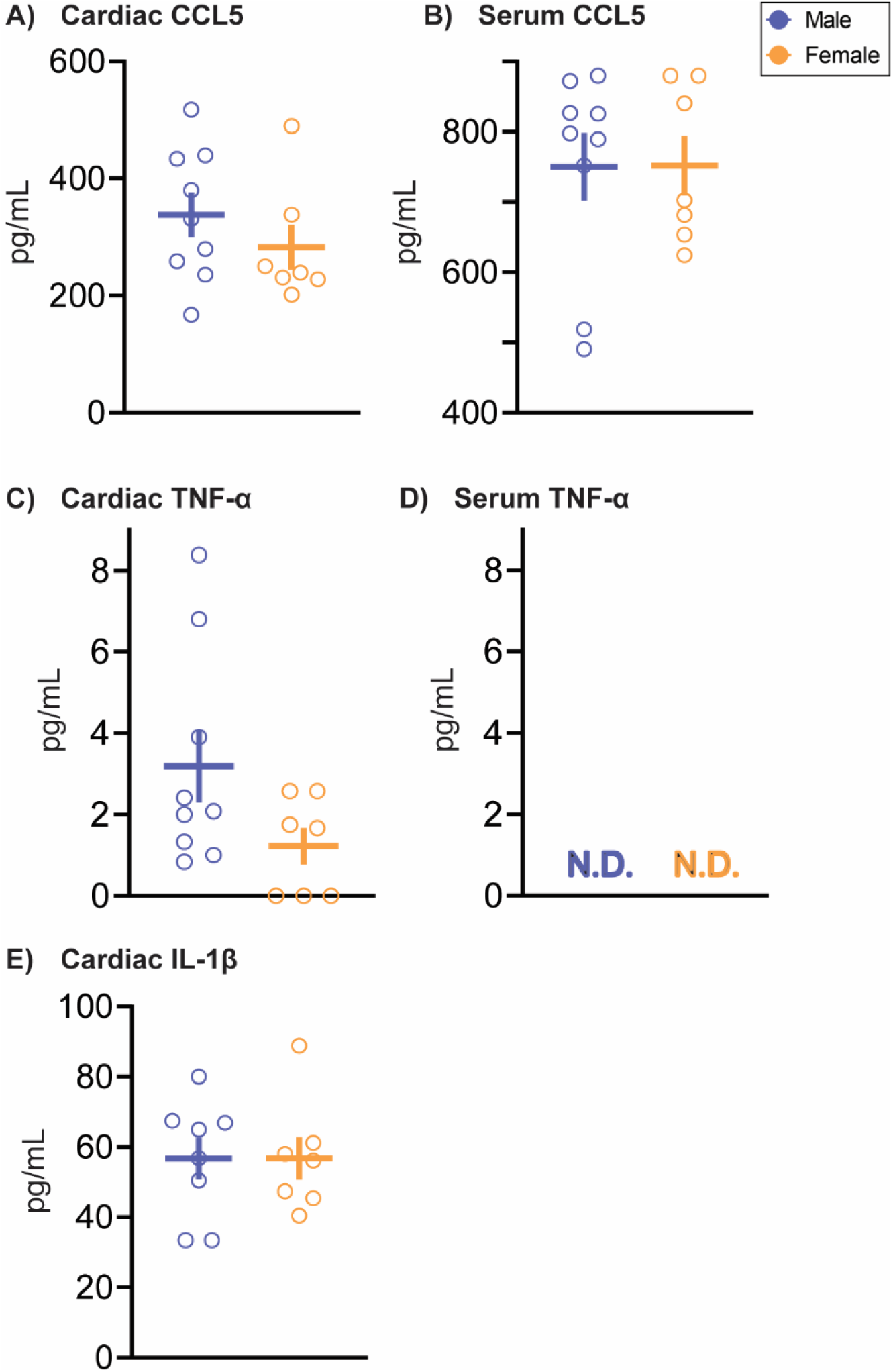
No sex differences are detected in naturally aged rats in protein expression of immune chemokines and cytokines CCL5, TNF-α, or IL-1β in heart and serum. At termination of all studies (∼13mo), rats were euthanized and heart and serum collected for ELISA analysis. There was no statistically significant sex difference in **(A-B)** CCL5 protein expression in heart or serum, **(C)** TNF-α expression in heart, or **(E)** IL-1β expression in heart. **(D)** There was no detectable (N.D.) TNF-α expression in serum in either sex. Protein concentration loaded for ELISAs was normalized using prior BCA analysis. Histograms depict average ± SEM. *n*=7-9 rats/sex/group.

**Supp. Fig. 3:**
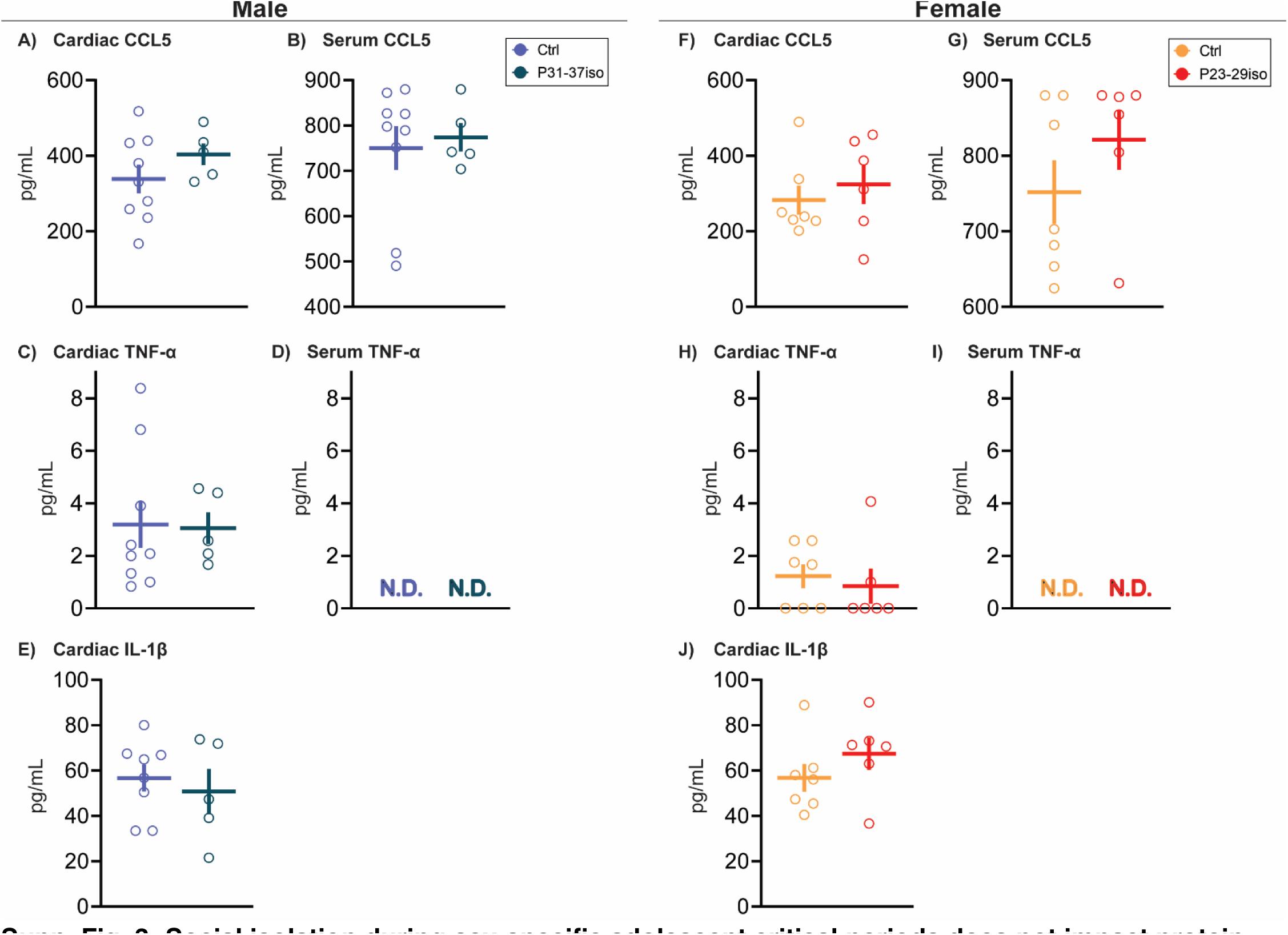
Social isolation during sex-specific adolescent critical periods does not impact protein expression of immune chemokines and cytokines CCL5, TNF-α, or IL-1β in heart and serum. Rats are single-housed during sex-specific critical periods for adolescent social development, P31-37 in males and P23-29 in females, and then re-housed with previous cage mates. At termination of all studies (∼13mo), rats were euthanized and heart and serum collected for ELISA analysis. There was no statistically significant effect of adolescent manipulation in either sex in **(A-B, F-G)** CCL5 protein expression in heart or serum, **(C, H)** TNF-α expression in heart, or **(E, J)** IL-1β expression in heart. **(D, I)** There was no detectable (N.D.) TNF-α expression in serum in either sex. Protein concentration loaded for ELISAs was normalized using prior BCA analysis. Histograms depict average ± SEM. *n*=5-9 rats/sex/group.

